# RNase L limits host and viral protein synthesis via inhibition of mRNA export

**DOI:** 10.1101/2021.04.18.440343

**Authors:** James M. Burke, Alison R. Gilchrist, Sara L. Sawyer, Roy Parker

## Abstract

RNase L is widely thought to limit viral protein synthesis by cleaving host rRNA and viral mRNA, resulting in translation arrest and viral mRNA degradation. Herein, we show that the mRNAs of dengue virus and influenza A virus largely escape RNase L-mediated mRNA decay, and this permits viral protein production. However, activation of RNase L arrests nuclear mRNA export, which strongly inhibits influenza A virus protein synthesis and reduces cytokine production. Importantly, the heterogeneous and temporal nature of the mRNA export block in individual cells permits sufficient production of antiviral cytokines from transcriptionally induced host mRNAs. This defines RNase L-mediated arrest of mRNA export as a key antiviral shutoff and cytokine regulatory pathway.

**One Sentence Summary:** RNase L-mediated shutoff of nuclear mRNA export limits viral protein synthesis and regulates antiviral cytokine production.

## Introduction

Double-stranded RNA (dsRNA) is a viral-associated molecular pattern that initiates innate immune programs in mammalian cells (*1, 2*). In response to dsRNA, cells induce the expression of antiviral proteins such as cytokines, which prime an antiviral state in infected and non-infected cells. Simultaneously, cells limit translation to reduce viral protein production. For decades, activation of the cytoplasmic endoribonuclease, RNase L, in response to viral double-stranded RNA (dsRNA) has been thought to limit viral protein synthesis by cleaving host ribosomal RNA (rRNA), resulting in arrest of translation (*3-7*). However, two recent studies indicate that RNase L-mediated cleavage of rRNA does not arrest translation. First, flaviviruses, including Zika virus and dengue virus (DENV), can replicate despite activating RNase L-mediated rRNA decay (*8*). Second, rapid and widespread decay of host mRNAs by RNase L accounts for RNase L-mediated reduction in translation (*9, 10*). In light of these findings, it is unclear how RNase L limits viral protein synthesis, how some viruses escape the effects of RNase L, and how RNase L activation affects host antiviral protein production.

## Results

### Dengue virus mRNAs escape RNase L-mediated mRNA decay and produce protein

DENV is a flavivirus (+ssRNA virus) that replicates in the cytoplasm. To investigate the mechanism by which DENV evades the effects of RNase L (*8*), we infected parental (WT) and RNase L-KO (RL-KO) A549 cells with dengue virus serotype 2 and performed single-molecule fluorescent *in situ* hybridization (smFISH) for *DENV* mRNA, *GAPDH* mRNA, which is degraded when RNase L is active (*9*), and also stained for DENV NS3 protein production by immunofluorescence.

DENV infected cells frequently activated RNase L, as observed by a strong reduction of *GAPDH* mRNA in most DENV-infected WT but not RL-KO cells (Fig. 1A,B and Extended Fig. S1). DENV-infected cells that activated RNase L showed a small, but not statistically significant, reduction in DENV mRNA levels as compared to WT cells that failed to activate RNase L, or RL-KO cells (Fig. 1C). Moreover, DENV protein production was strongly correlated with DENV mRNA levels (Fig. 1D), and only showed small decreases in cells with activated RNase L (Fig. 1E). Thus, DENV RNA is largely unaffected by RNase L-mediated mRNA decay. Consequently, this permits DENV protein production, consistent with cytoplasmic mRNAs having the capacity to be translated during the RNase L response (*9, 10*).

**Fig. 1.**
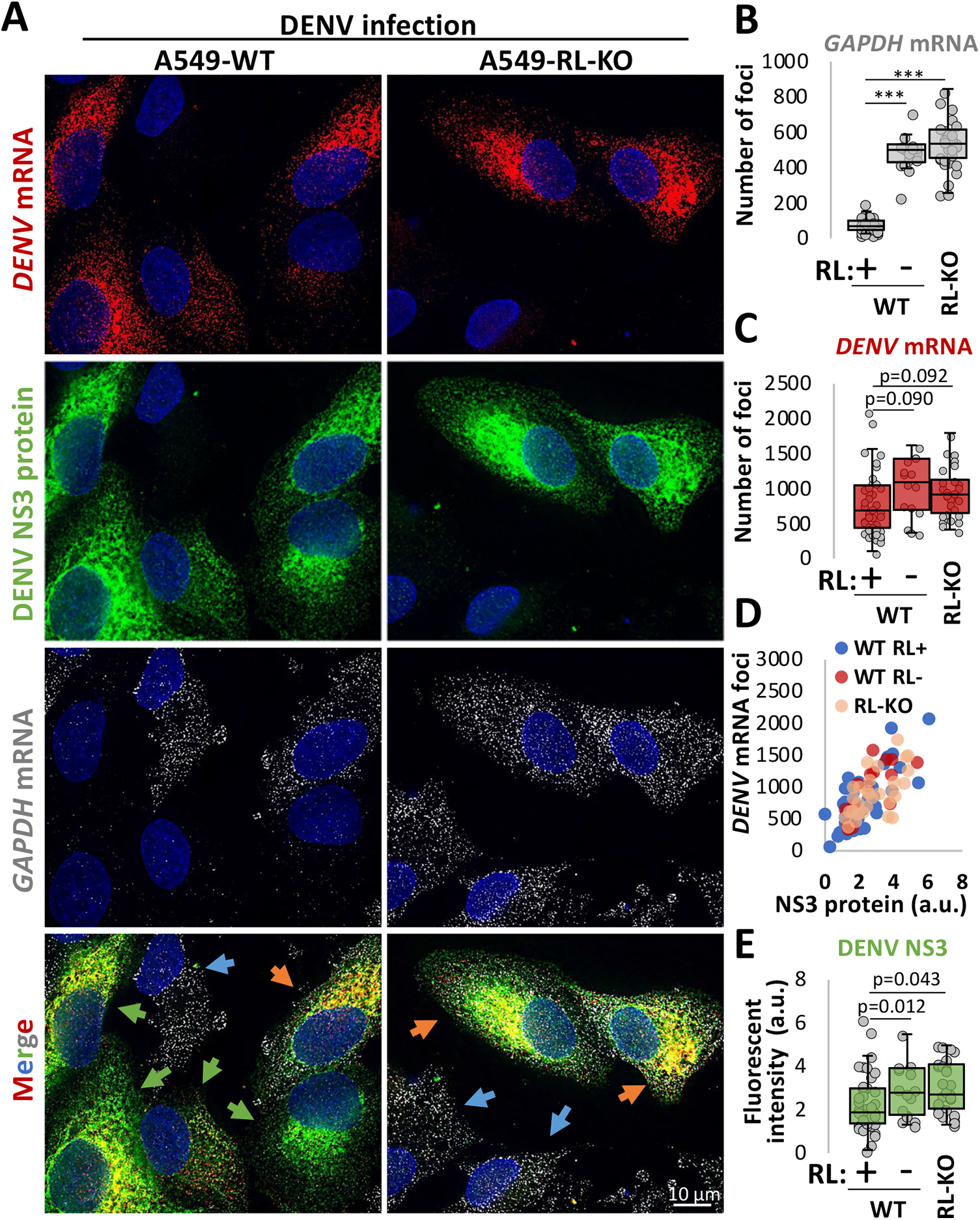
Dengue virus mRNAs escape RNase L-mediated mRNA decay and are translated. (**A**) smFISH for *DENV* mRNA and *GAPDH* mRNA and immunofluorescence for DENV NS3 protein in WT and RL-KO A549 cells forty-eight hours post-infection with DENV (MOI=0.1). Images are from a single z-plane. Orange arrow: RNase L not activated (*GAPDH* mRNA not degraded); green arrow: RNase L activated (*GAPDH* mRNA degraded); Blue arrow: Uninfected cells (neither DENV mRNA nor NS3 detected). (**B**) Quantification of GAPDH mRNA in WT and RL-KO cells. WT cells in which *GAPDH* mRNA levels were lower than the lowest level observed in RL-KO cells were designated as RNase L active (RL+). WT cells with GAPDH mRNA levels above that threshold were designated RNase L inactive (RL-). (**C**) Quantification of *DENV* mRNA foci. (**D**) Scatterplot of DENV mRNA foci and DENV NS3 immunofluorescence. (**E**) similar to (**B** and **C**) but quantifying the integrated density (arbitrary units: a.u.) of DENV NS3 immunofluorescence as represented in (**A**).

### The host immune response inhibits mRNA export via RNase L activation

Examination of host cytokine mRNAs induced by DENV infection in WT and RL-KO A549 cells revealed two important points. First, we observed that antiviral cytokine mRNAs largely escape RNase L-mediated mRNA decay. This is based on the observation that in some WT cells that activated RNase L-mediated mRNA decay (degraded GADPH mRNA), *IFN-λ1* and *IFN**-**β* mRNAs were abundant in the cytoplasm, often at levels comparable to those observed in RL-KO cells (Fig. 2A,B and Extended Figs S2A,B and S3A,B). Moreover, while median GAPDH mRNA levels were reduced more than tenfold by RNase L (Fig. 1B), median *IFN* mRNA levels were only reduced twofold by RNase L (Fig. 2C and Extended Fig. S3C). Thus, cytokine mRNAs that reach the cytoplasm largely escape RNase L-mediated mRNA decay during DENV infection, consistent with our observations during the dsRNA response (*9*).

**Fig. 2.**
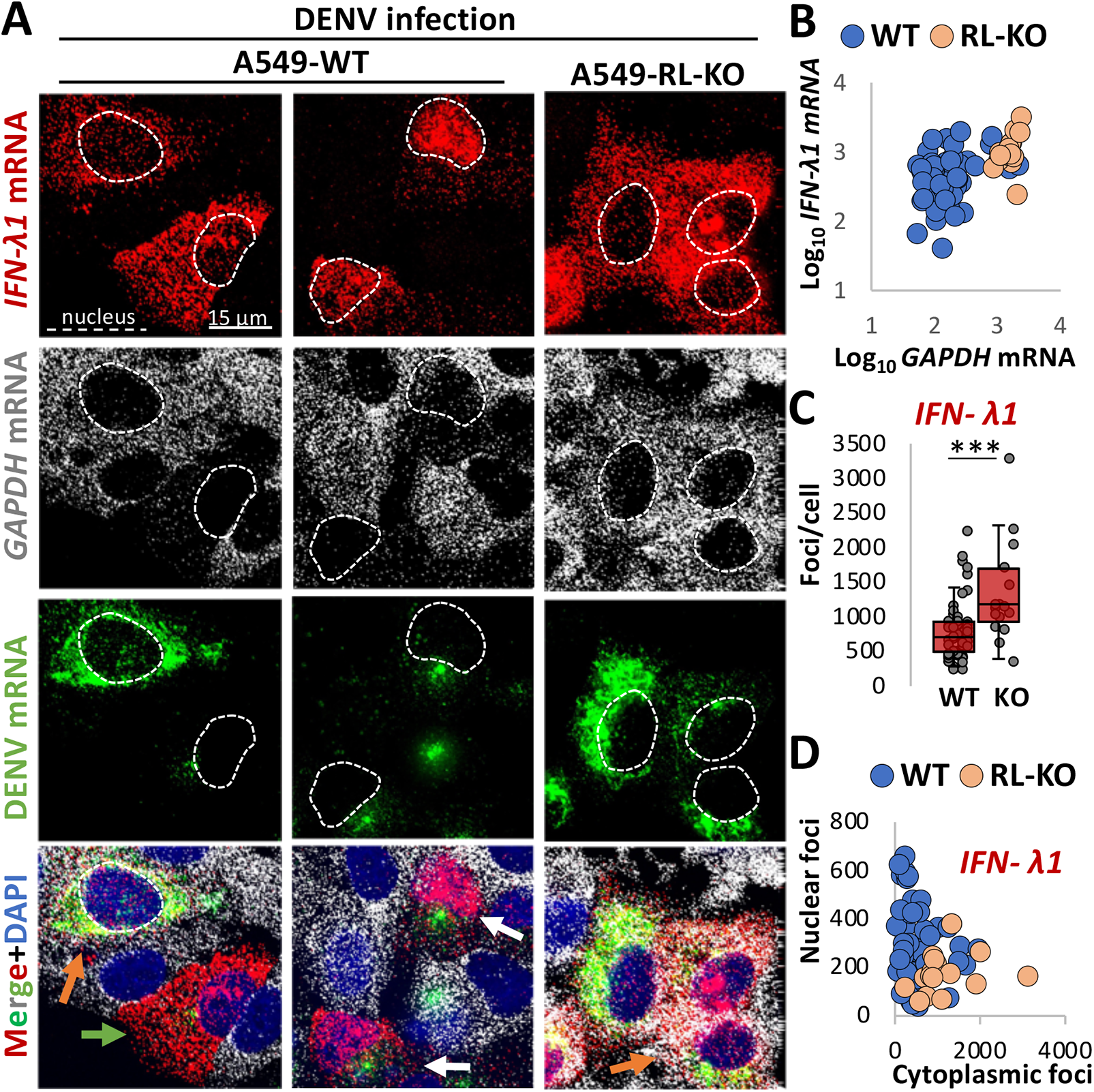
RNase L inhibits nuclear export of interferon mRNAs in response to DENV infection. **(A)** smFISH for *IFN-λ1* mRNA, *GAPDH* mRNA, and *DENV* mRNA in WT and RL-KO A549 cells forty-eight hours post-infection with DENV (MOI=0.1). Orange arrow: DENV-positive but RNase L not activated (*GAPDH* mRNA not degraded); green arrow: DENV positive, RNase L activated (*GAPDH* mRNA degraded), and *IFN-λ1* mRNA localized to cytoplasm; white arrow: DENV-positive, RNase L activated, and *IFN-λ1* mRNA retained in the nucleus. **(B)** Scatter plot of *GAPDH* mRNA and *IFN-λ1* mRNA in individual cells. **(C)** Quantification of smFISH foci in WT and RL-KO cells infected with DENV and that induced *IFN-λ1* mRNA expression. (**D**) Scatter plot of the number of smFISH foci of *IFN-λ1* mRNA localized to the nucleus or cytoplasm in individual WT and RL-KO cells as represented in (**A**).

Surprisingly, we also observed accumulation of *IFN-β* and *IFN-λ1* mRNAs within the nucleus of some DENV-infected WT cells (Fig. 2A,D and Extended Figs. S2A,B and S3A-D). In principle, this mRNA export block could be mediated by DENV or be consequence of the cellular response to dsRNA. To distinguish these possibilities, we transfected cells with poly(I:C), a viral dsRNA mimic, and examined if mRNA export was affected. We observed that poly(I:C) transfection was sufficient to activate RNase L, to induce *IFN* mRNAs, and to block mRNA export in a fraction of WT cells (Fig. 3A-C and Extended Fig. 4A-C). This demonstrates a new aspect of the host innate immune response that blocks nuclear mRNA export.

**Fig. 3.**
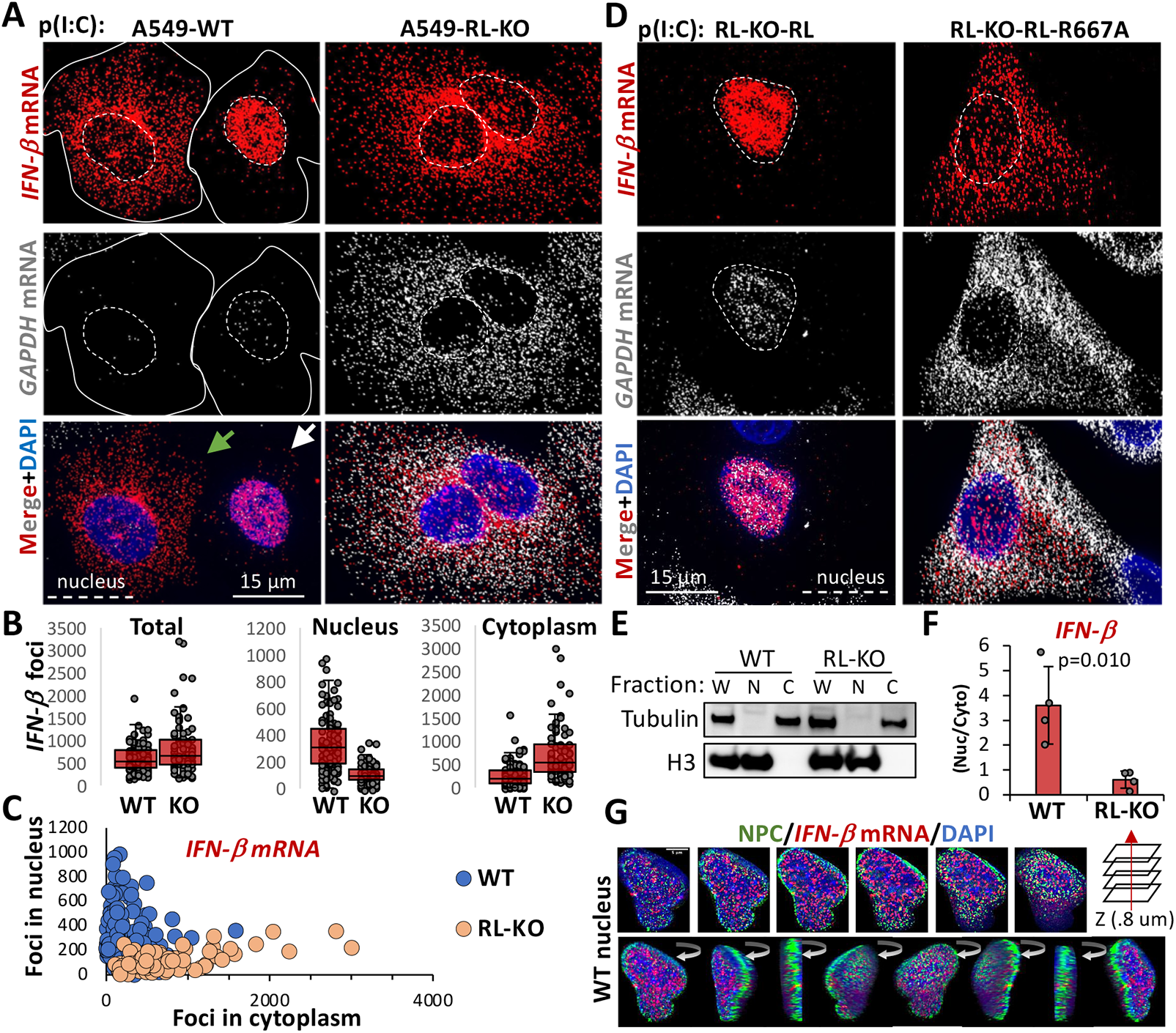
RNase L inhibits nuclear export of interferon mRNAs in response to dsRNA. **(A)** smFISH for *IFN-β* and *GAPDH* mRNAs in wild-type (WT) and RNase L-KO (RL-KO) A549 cells eight-hours post-poly(I:C) lipofection. Solid lines mark the cell boundary of WT cells since it is difficult to view cell boundaries with nuclear-retained IFN mRNA. (**B**) Quantification of *IFN-β* smFISH foci, as represented in (A), in individual cells (dots). **(C)** Scatter plot of the nuclear and cytoplasmic *IFN-β* foci in individual WT and RL-KO cells, as represented in (**A**). (**D**) smFISH for *IFN-β* and *GAPDH* mRNAs in RL-KO cells expressing RNase L or RNase L-R667A catalytic mutant via lentiviral transduction post-poly(I:C) lipofection. (**E**) Immunoblot of whole cell (W), nuclear (N), and cytoplasmic (C) fractions from WT and RL-KO cells six hours post poly(I:C) used for RT-qPCR. (**F)** qRT-PCR quantification of *IFN-β* mRNA from nuclear-cytoplasmic fractions as shown in (**E**). Bars represent the average 2^(-*Δ*Ct [nuclear-cytoplasm]) +/-S.D (n=4). Dots represent independent experiments. The p-value was determined using student’s ttest. **(G)** Immunofluorescence for nuclear pore complex (NPC) and smFISH for *IFN-β* mRNA in a WT cell displaying nuclear retention of *IFN-β* mRNA.

Several observations demonstrate that this block to mRNA export requires activation of RNase L-mediated mRNA degradation. First, nuclear retention of *IFN* mRNAs was only observed in DENV-infected WT cells that activated RNase L-mediated decay of *GAPDH* mRNA (Fig. 2A,D). In contrast, neither WT cells that failed to activate RNase L (as assessed by the lack of *GAPDH* mRNA degradation) nor RL-KO cells that induced cytokine mRNAs displayed the mRNA export block (Fig. 2A,D). Second, the mRNA export block in response to poly(I:C) was also RNase L-dependent, as we did not observe *IFN* mRNA accumulation in the nucleus of RL-KO cells (Fig. 3A-C and Extended Fig. S4A-C). Third, the block to mRNA export requires RNase L catalytic activity since rescuing the RL-KO cells with RNase L, but not RNase L-R667A (a catalytic mutant), restored nuclear retention of *IFN**-**β* mRNA in response to poly(I:C) (Fig. 3D). Nuclear accumulation of IFN**-***β* mRNAs was also observed by RT-qPCR of nuclear and cytoplasmic biochemical fractions (Fig. 3E,F). Thus, activation of RNase L is sufficient to induce a block to nuclear mRNA export.

Immunofluorescence assay showed that nuclear-retained *IFN**-**β* mRNA is internal to the nuclear pore complex (NPC) and not retained at the site of transcription (Fig. 3G), which can be observed with many RNA processing defects (*11*). The block to nuclear export is independent of PKR, since nuclear accumulation of mRNAs after poly(I:C) treatment is also observed in PKR-KO cells (Extended Fig. S4D). The nuclear block appears independent of RNase L-mediated apoptosis since the VZAD caspase inhibitor did not inhibit it (Extended Fig. S4E).

Our observations indicate that RNase L activation inhibits the CRM-1 mRNA export pathway, which is a specialized mRNA export pathway used by mRNAs without introns and/or that contain AU-rich elements (AREs) (*12*). The *IFN-β* mRNA, which lacks introns and contains AREs, is thought to exit the nucleus by the CRM-1 mRNA export pathway (*13*). Indeed, treatment of RL-KO cells with the CRM-1 inhibitor, leptomycin B, phenocopied the RNase L-dependent nuclear retention of *IFN-β* mRNA observed in WT cells (Extended Fig. S4F). Thus, nuclear accumulation of *IFN-β* mRNA is a result of RNase L-dependent inhibition of CRM-1 mediated mRNA export.

RNase L activation also inhibits the NFX1-TREX bulk mRNA export pathway that acts on spliced mRNAs since *IFN-λ1* mRNA contains introns and is insensitive to leptomycin B (Extended Fig. S4G). Moreover, the GAPDH mRNA, which is also spliced and presumably exported by the NFX-TREX pathway, also shows nuclear retention in cells with activated RNase L (Fig. 3D). These observations argue RNase L activation inhibits both the CRM-1 and the NFX1-TREX mRNA export pathways.

### Activation of RNase L inhibits influenza virus mRNA export and protein synthesis

We hypothesized that RNase L-mediated inhibition of mRNA export may be important for limiting viral protein synthesis for viruses that must export their mRNAs from the nucleus. For example, influenza (-ssRNA virus) replicates in the nucleus and must export its mRNAs to the cytosol and is inhibited 100-to 1000-fold by RNase L (*14*). To test if RNase L activation blocked influenza mRNA export, we infected WT and RL-KO A549 cells with influenza A/Udorn/72 virus (IAV). Since the IAV NS1 protein strongly attenuates RNase L activation (*14*) (Extended Fig. S5A), we transfected cells with poly(I:C) one-hour post-infection to promote RNase L activation. We performed smFISH seven hours post-infection, when influenza A/Udorn/72 viral output is in log phase growth and when RNase L reduces viral output by greater than 100-fold (*14*).

An important result was that most WT cells that activated RNase L (as assessed by degradation of the *GAPDH* mRNA) limited the export of IAV *NA* and *NS1* mRNAs (Fig. 4A,B and Extended Fig. S5B-E). Nuclear retention of these mRNAs did not occur in WT cells that failed to activate RNase L-mediated mRNA decay nor in RL-KO cells. This demonstrates that RNase L activation leads to the inhibition of IAV mRNA export.

**Fig. 4.**
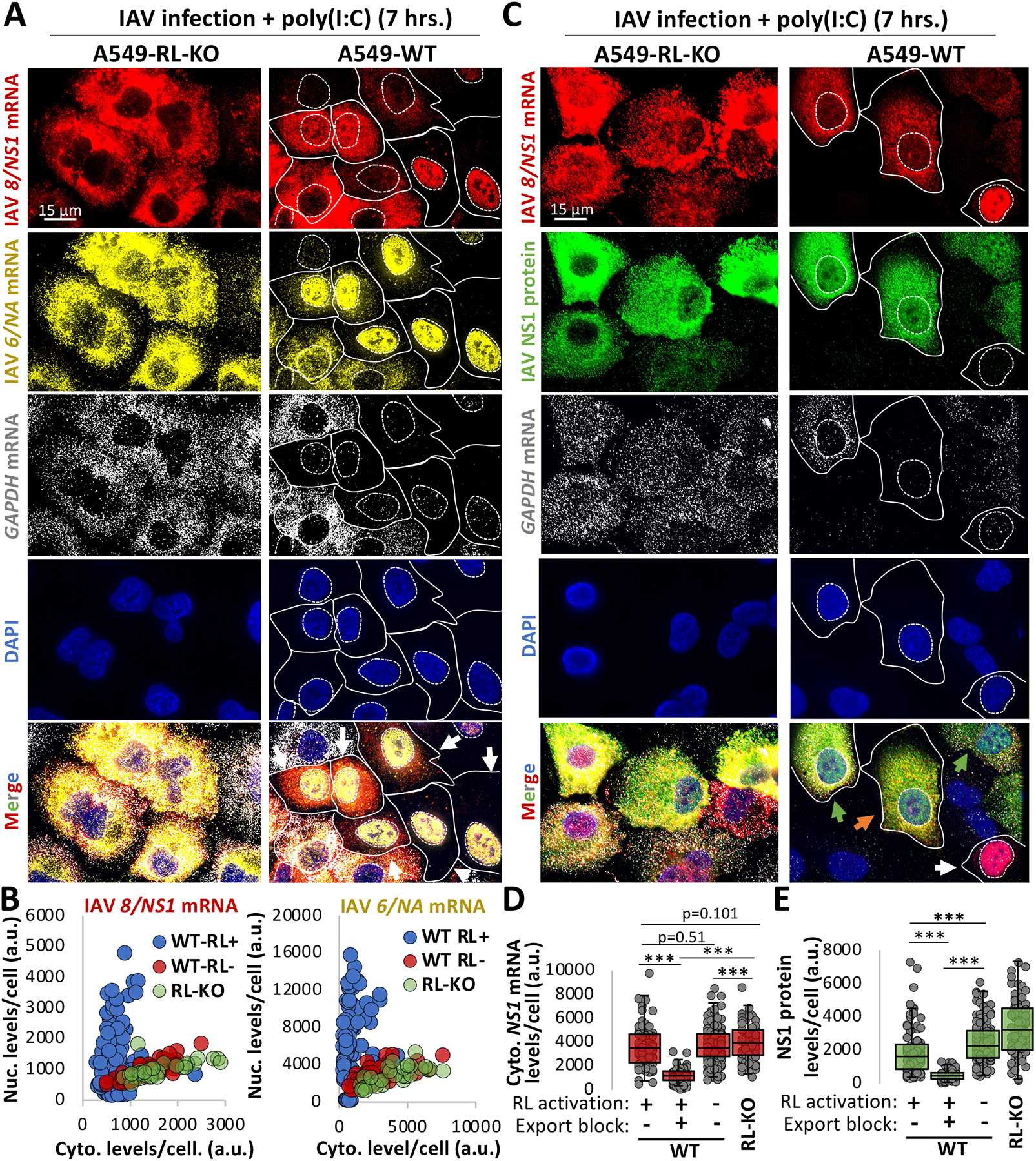
RNase L-mediated inhibition of mRNA export inhibits influenza virus protein synthesis. (**A**) smFISH for IAV *NA* and *NS1* mRNAs seven-hours post-infection (MOI=0.5). Cells were transfected with poly(I:C) one-hour post-infection. *GAPDH* mRNA degradation is a marker for RNase L activation. Additional images are shown in Extended Fig. 5B,C. White arrows (nuclear retention of *NA* and *NS1*); blue arrows (nuclear retention of *NA* but not *NS1*). (**B**) scatter plot of nuclear intensity and cytoplasmic intensity (arbitrary units: a.u.) of *NA* or *NS1* mRNAs from individual WT cells with active RNase L (RL+; loss of *GAPDH* mRNA) or inactive RNase L (RL-; contain *GAPDH* mRNA) and RL-KO cells. (**C**) similar to (**A**) but smFISH for *NS1* mRNA and *GAPDH* mRNA and immunofluorescence staining of NS1 protein. Green arrow (RL-, cytoplasmic *NS1* mRNA+); orange arrow (RL+, cytoplasmic *NS1* mRNA+); white arrow (RL+, nuclear *NS1* mRNA retention+). (**D**) cytoplasmic staining intensity of *NS1* mRNA in RL-WT cells, RL+ WT cells displaying either *NS1* mRNA nuclear retention (export block +) or cytoplasmic localization (export block-), and RL-KO cells. WT cells with export block were defined as having a nuclear to cytoplasmic *8/NS1 mRNA* intensity ratio that was higher than the maximum ratio observed in RL-KO cells for each replicate (Extended Fig. 6C). (**E**) similar to (**D**) but plotting immunofluorescent intensity of NS1 protein. Statistical significance (*p<0.05; **p<0.005; ***p<0.0005) was determined by ttest analysis.

We observed that in cells that activated RNase L-mediated mRNA decay, but failed to trigger the mRNA export block, influenza mRNAs were abundant in the cytoplasm (Fig. 4A,C and Extended Figs. S5B,F and S6A), at levels comparable to those observed in WT cells that did not activate RNase L and RL-KO cells (Fig. 4D and Extended Figs. S5B-F). This argues that IAV mRNAs escape RNase L-mediated mRNA decay and could potentially be translated for protein production, similar to DENV mRNAs (Fig. 1). This suggested that the RNase L-mediated mRNA export block was the mechanism by which RNase L limits influenza protein synthesis.

To test this hypothesis, we examined the relationship between influenza NS1 protein production and the *NS1* mRNA export block in individual cells (Fig. 4C and Extended Fig. S6A,B,C). We observed that cells displaying RNase L activation (*GADPH* mRNA degradation), but successful *NS1* mRNA export, produced NS1 protein at levels comparable, albeit slightly less, to WT cells that did not activate RNase L and RL-KO cells (Fig. 4C-E). In contrast, WT cells in which *NS1* mRNA export was inhibited by RNase L activation displayed significantly less NS1 protein production (Fig. 4C-E). Lastly, cytoplasmic levels of *NS1* mRNA strongly correlated with NS1 protein levels with or without RNase L activation (Extended Fig. S6B). These observations demonstrate that RNase L-mediated inhibition of mRNA export is a primary mechanism by which RNase L reduces influenza protein production.

### RNase L-mediated inhibition of mRNA export limits antiviral protein production

RNase L-mediated inhibition of mRNA export would also be expected to limit translation of transcriptionally induced antiviral mRNAs, which could be detrimental to the antiviral response. However, since this response was heterogeneous with respect to individual cells (Figs. 2 and 3), we suspected that the RNase L-mediated mRNA export block would reduce, but not abolish, antiviral cytokine production. Such a function would be important for ensuring cytokine production while potentially preventing the overproduction of cytokines, which can cause cytokine storm phenomena.

Three primary observations support these hypotheses. First, RNase L expression reduced, but did not abolish, secretion of IFN-*β* and IFN-λ1 proteins in manner dependent on its catalytic activity in response to poly(I:C) (Fig. 5A and Extended Fig. S7), consistent with a previous study (*15*).

**Fig. 5.**
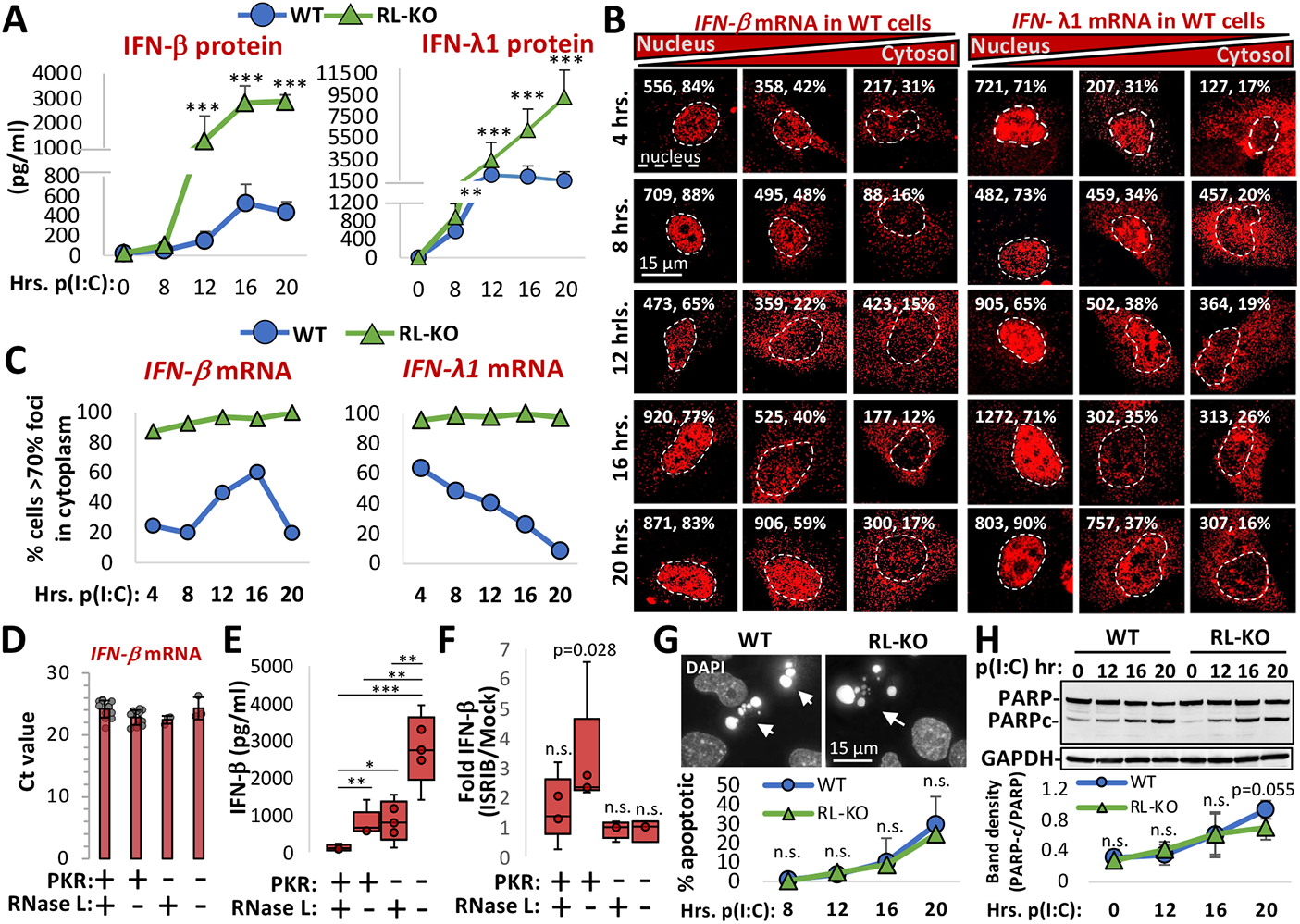
RNase L-mediated mRNA export block contributes to inhibition of antiviral cytokine production. (**A**) ELISAs for IFN-*β* and IFN-λ1. Markers represent the mean S.D. from at least six replicates. Statistical significance (*p<0.05; **p<0.005; ***p<0.0005) was determined by ttest analysis. (**B**) WT A549 cells displaying varying degrees of nuclear retention of *IFN-β* or *IFN-λ1* mRNA post-poly(I:C). The number and percentage of smFISH foci localized to the nucleus are shown in the top right corners. Nuclei were determined by DAPI staining (not shown for space and clarity) and are outlined. Additional images are shown in Extended Figs. S8 and S9. (**C**) The percentage of cells displaying 70% or greater *IFN-β* or *IFN-λ1* smFISH foci in the cytoplasm derived from Extended Fig. S9D. (**D**) RT-qPCR for *IFN-β* mRNA in WT, RL-KO, PKR-KO, and RL/PKR-dKO cell lines six hours post-poly(I:C). Bars represent the average cycle threshold (Ct) +/-S.D. Each dot represents a Ct value from an individual experiment. (**E**) Quantification of IFN-*β* via ELISA twelve hours post-poly(I:C). (**F**) Fold change in IFN-*β* secretion via ELISA twelve hours post-poly(I:C) with or without co-treatment with ISRIB. (**G**) WT and RL-KO cells stained with DAPI to visualize apoptotic cells. The graph below shows the mean percentage and S.D. of apoptotic cells. (**H**) Immunoblot analysis of PARP and caspase-mediated cleavage fragment of PARP (PARPc). The graph below shows the mean +/- S.D. of PARPc/PARP ratio.

Second, nuclear retention of *IFN-β* and *IFN-λ1* mRNAs was differential with respect to time (Fig. 5B,C and Extended Figs. S8A-D and S9A-D), and this correlated with RNase L-dependent repression of protein production (Fig. 5A). Specifically, ∼75% of WT cells displayed nuclear retention of *IFN-β* mRNA prior to twelve hours post-poly(I:C) (Fig. 5C), when RNase L inhibited IFN-*β* levels by tenfold (Fig. 5A). At twelve-and sixteen-hours post-poly(I:C), cytoplasmic *IFN-β* mRNA levels increased in WT cells (Fig. 5C), which preceded/coincided with an increase in IFN-*β* (Fig. 5A). In contrast, fewer cells (∼35%) displayed nuclear retention of *IFN-λ1* mRNA at early times post-poly(I:C) (Fig. 5C), and this correlated with comparable IFN-λ1 levels from WT and RL-KO cells at eight- and twelve-hours post-poly(I:C) (Fig. 5A). However, localization of the *IFN-λ1* mRNA progressively increased to nucleus over time in WT cells (Fig. 5C), correlating with a cessation of IFN-λ1 secretion at late times post-poly(I:C) (Fig. 5A). These data show that RNase L-mediated inhibition of *IFN* mRNA nuclear export correlates with reduction in IFN protein production.

Third, several observations indicate that RNase L inhibits IFN-*β* protein production at twelve hours post-poly(I:C) specifically via the export block as opposed to mRNA decay. The twofold reduction of *IFN-β* mRNA by RNase L prior to twelve hours post-poly(I:C) is insufficient to account for the tenfold reduction of IFN-*β* protein observed at twelve hours post-poly(I:C) (Fig. 5A,D and Extended Fig. S9C). Moreover, despite an equivalent ∼twofold reduction in *IFN-λ1* and *IFN-β* mRNAs by RNase L during this time, IFN-λ1 protein levels were largely unaffected by RNase L (Fig. 5A and Extended Fig. S9C,D), correlating with *IFN-λ1* mRNA being predominantly cytoplasmic whereas *IFN-β* mRNA was primarily nuclear at early times post-poly(I:C) (Fig. 5C). These data argue that nuclear retention, as opposed to RNA decay, of the *IFN-β* mRNA accounts for reduced IFN-*β* protein production.

We also considered the possibility that RNase L limits IFN-*β* protein production by activating eIF2*α* kinases (*9*), in particular PKR (*16*), which would promote phosphorylation of eIF2*α* (p-eIF2*α*) and repress translation. However, the cumulative increase of IFN-*β*production in RNase L/PKR double knockout cells in comparison to PKR-KO or RL-KO cells argues that RNase L inhibits IFN-*β* production independently of PKR (Fig. 5E). We note that neither PKR nor RNase L markedly affected *IFN-β* mRNA levels (Fig. 5D). Moreover, ISRIB treatment, which counteracts the effects of p-eIF2*α* (*17*), failed to increase IFN-*β* in cells that express RNase L (Fig. 5F). Thus, RNase L reduces translation of *IFN-β* mRNA independently of p-eIF2*α*.

Lastly, RNase L-mediated apoptosis (*18, 19*) is not responsible for limiting IFN-*β* production at twelve hours post-poly(I:C), since we did not observe a notable number of apoptotic cells nor caspase-mediated PARP cleavage prior to twelve hours post-poly(I:C) (Fig. 5G,H), consistent with live-cell imaging studies (*20*). Moreover, we did not observe a substantial difference in the rate of cells initiating apoptosis nor in the rate of caspase-mediated PARP cleavage between WT and RL-KO cells (Fig. 5G,H).

Taken together, these observations argue that global reductions in translation via mRNA decay, translation arrest, or apoptosis do not account for RNase L-mediated inhibition of IFN-*β*. Instead, our data demonstrate that RNase L reduces type I and type III IFN production by blocking the export of their mRNAs to the cytoplasm.

## Discussion

We present several observations documenting new aspects of the innate immune response. First, we show that cytoplasmic DENV and IAV mRNAs escape RNase L-mediated decay and, consequently, produce protein during the RNase L response (Figs. 1 and 4). These data indicate that RNase L activation does not arrest translation via rRNA cleavage, consistent with recent studies (*9, 10*), and elucidate how some viruses, such as ZIKA virus, can synthesize proteins despite robust RNase L-mediated rRNA decay (*8*). Similar to host antiviral mRNAs (*9, 10*), the high transcriptional rates of viral mRNAs and their structure, as well as their localization to replication factories and association with viral proteins (*8*), likely contributes to their ability to escape the effects of RNase L-mediated mRNA decay.

We also identified a new mechanism by which RNase L reduces host and viral gene expression, whereby RNase L activation inhibits mRNA export of host and viral mRNAs, which reduces their translation by preventing their association with ribosomes in the cytoplasm. This is based on the observations that host and viral mRNAs accumulate in the nucleus of cells that have activated RNase L, and this correlates to reduced protein expression from these transcripts (Figs. 2-5).

The mechanism by which activation of RNase L inhibits mRNA export remains to be discovered. Several observations are consistent with RNA export being inhibited following widespread cytoplasmic RNA decay due to an influx of cytoplasmic RNA-binding proteins (RBPs), which then limit the interaction of RNA export factors with nuclear mRNAs. For example, in addition to RNase L-mediated mRNA decay, widespread degradation of host cytoplasmic mRNAs by the SOX and VHS nucleases encoded by Kaposi’s sarcoma-associated herpesvirus (KSHV) and herpes simplex virus 1 (HSV-1), respectively, leads to the import of cytosolic RBPs into the nucleus in conjunction with mRNA export inhibition (*9, 20-24*). Further supporting this model, RNase L-mediated inhibition of mRNA export is dependent on its catalytic activity and occurs after mRNA decay (Fig. 3A-D). Elucidating the mechanism by which RNase L-mediated mRNA decay inhibits mRNA export will be a key focus of future studies.

One function of RNase L-mediated inhibition of mRNA export is to limit viral protein production by limiting gene expression from viruses that replicate in the nucleus. Indeed, our data suggest that this is a primary mechanism by which RNase L limits influenza protein production since nuclear accumulation of influenza mRNAs is required for a reduction in their translation in cells with activated RNase L (Fig. 4). Since most DNA viruses must export their mRNAs from the nucleus and can activate RNase L (*25-28*), this could be a broad antiviral mechanism.

A second function of RNase L-mediated inhibition of mRNA export is to regulate cytokine production. We observed that the reduction of IFN-*β* protein due to RNase L correlates with the mRNA export block and cannot be explained by RNase L-mediated global reduction in translation, mRNA degradation, or enhanced apoptosis (Fig. 5). Thus, RNase L activation limits cytokine production by trapping cytokine mRNAs in the nucleus in a fraction of the cells (Fig. 5). We suggest that the heterogenous and temporal nature between individual cells allows for sufficient production of cytokines to limit viral replication (Figs. 3A,B and 5A-C and Extended Figs. S7-9) (*29)*. Moreover, our data indicate that type III IFNs are less repressed by RNase L than type I IFNs at early times post-dsRNA (Fig 5A-C and Extended Figs. 7-9). This would promote localized epithelial cell-specific antiviral signaling prior to systemic type I IFN signaling (*30*). Combined, these functions may prevent systematic overproduction of cytokines, which can lead to cytokine storm, sepsis, and autoimmune disorders in response to viral infection (*31-33*).

## Materials and Methods

### Cell culture

The A549 cell line was provided by Dr. Christopher Sullivan (The University of Texas at Austin). The A549 RNase L knockout (RL-KO) cell line and RNase L and RNase L-R667A RL-KO cell lines are described in (*9*). The generation of PKR-KO and RL/PKR-KO and lines are described in (*20*). Cells were maintained at 5% CO2 and 37 degrees Celsius in Dulbecco’s modified eagle’ medium (DMEM) supplemented with fetal bovine serum (FBS; 10% v/v) and penicillin/streptomycin (1% v/v). Cells tested negative for mycoplasma contamination by the BioFrontiers cell culture core facility. Cells were transfected with poly(I:C) HMW (InvivoGen: tlrl-pic) using 3-μl of lipofectamine 2000 (Thermo Fisher Scientific) per 1-ug or poly(I:C). Cells were treated with Caspase Inhibitor Z-VAD-FMK (Promega: G7231) at 20μM concentration.

### Virus infections

A549 cells were infected with dengue virus serotype 2 16681 strain at an MOI of 0.1. Cells were fixed 48 hours post-infection. Cells were infected with influenza A/Udorn/72 virus strain at MOI of 0.5. Cells were fixed 7 hours post-infection.

### Immunoblot analyses

Immunoblot analysis was performed as describe in (*9*). Rabbit anti-GAPDH (Cell Signaling Technology: 2118L) was used at 1:2000. Anti-rabbit IgG, HRP-linked antibody (Cell Signaling Technology: 7074S) was used at 1:3000. Anti-mouse IgG, HRP-linked antibody (Cell Signaling Technology: 7076S) was used at 1:10,000. Rabbit anti-PARP (Cell Signaling Technology: 9452S) was used at 1:1,500. Histone H3 antibody (Fisher Scientific; NB500-171) was used at 1:1000.

### Immunofluorescence and single-molecule FISH

smFISH was performed following manufacturer’s protocol (https://biosearchassets.blob.core.windows.net/assets/bti_custom_stellaris_immunofluorescence_seq_protocol.pdf). GAPDH smFISH probes labeled with Quasar 570 Dye (SMF-2026-1) or Quasar 670 Dye (SMF-2019-1) were purchased from Stellaris. Custom IFNB1, IFNL1, IAV, and DENV smFISH probes were designed using Stellaris smFISH probe designer (Biosearch Technologies) available online at http://www.biosearchtech.com/stellaris-designer. Reverse complement DNA oligos were purchased from IDT (Extended data file 1). The probes were labeled with Atto-633 using ddUTP-Atto633 (Axxora: JBS-NU-1619-633), with ATTO-550 using 5-Propargylamino-ddUTP (Axxora; JBS-NU-1619-550), or ATTO-488 using 5-Propargylamino-ddUTP (Axxora; JBS-NU-1619-488) with terminal deoxynucleotidyl transferase (Thermo Fisher Scientific: EP0161) as described in (*34*).

For immunofluorescence detection of IAV NS1, cells were incubated with the anti-Influenza A virus NS1 rabbit antibody (GeneTex; GTX125990) at 1:1000 for two hours, washed three times, and then incubated with the Goat Anti-Rabbit IgG H&L (FITC) (Abcam; ab6717) at 1:2000 for one hour. After three washes, cells were fixed and then smFISH protocol was performed. For detection of DENV NS3, cells were incubated with anti-Dengue virus NS3 protein antibody (GeneTex: GTX124252) at 1:1000 for two hours, and then incubated with Goat Anti-Rabbit IgG H&L (Alexa Fluor® 647) (Abcam: ab150079) for one-hour. Mouse anti-Nuclear Pore Complex Proteins antibody (Abcam; ab24609) was used at 1:500 and detected with goat anti-mouse IgG H&L (FITC) (Abcam; ab97022) used at 1:2000.

### Microscopy and Image Analysis

Cover slips were mounted on slides with VECTASHIELD Antifade Mounting Medium with DAPI (Vector Laboratories; H-1200). Images were obtained using a wide field DeltaVision Elite microscope with a 100X objective using a PCO Edge sCMOS camera. Between 10-15 Z planes at um/section were taken for each image. Deconvoluted images were processed using ImageJ with FIJI plugin. Z-planes were stacked and minimum and maximum display values were set in ImageJ for each channel to properly view fluorescence. Fluorescent intensity was measured in Image J. Single cells were outlined by determining the cell boundaries via background fluorescence and mean intensity and integrated intensity were measured in the relevant channels. Imaris Image Analysis Software (Bitplane) (University of Colorado-Boulder, BioFrontiers Advanced Light Microscopy Core) was used to quantify smFISH foci in nucleus and cytoplasm. Single cells were isolated for analysis by defining their borders via background fluorescence. Total foci above background threshold intensity were counted. Afterwards, the nucleus marked with DAPI was masked, and foci were counted in the cell at the same intensity threshold cut-off, yielding the cytoplasmic foci count, from which the nuclear foci number could be determined.

### RT-qPCR

RT-qPCR for IFN-B1 was performed as described in (*9*). WT and RL-KO A549 cells (12-well; 60% confluent) were transfected with or without poly(I:C). Six hours post-transfection, crude nuclear and cytosolic fractions were obtained as described in (*35*). RNA was extracted from each fraction, treated with DNase I (NEB) for 15 minutes, and re-purified via ethanol (75%) sodium acetate (0.3M) precipitation, and re-suspended in 15-ul of water. Equal volumes of RNA from each fraction was then reverse transcribed using super script III reverse transcriptase (Thermo Fisher Scientific) and polydT(_20_) primer (Integrated DNA Technologies). cDNA was diluted to 100-ul. 2-ul of cDNA was added to qPCR reaction containing iQ SYBR green master mix (Bio-Rad) and 10-pmol of gene-specific primers. Reactions were run in duplicate or triplicate (technical replicates) on CFX96 qPCR machine (Bio-Rad) using standard two-step cycle.

### Quantification of secreted cytokines

WT and RL-KO A549 cells (6-well format, 1 ml or medium, 70% confluency) were transfected with poly(I:C). At six- and twelve-hours post-poly(I:C), 50-ul of medium was removed from well and immediately assayed via ELISA using manufacturer’s instructions. IFN beta human ELISA kit (Thermo Fisher Scientific; 414101) was used to quantify IFNB1. Human IL-29/IFN-lambda 1 ELISA Kit (Novus Biologicals; NBP1-84819) was used to quantify IFNL1. Time zero was taken by removing 50-ul of medium prior to poly(I:C) transfection.

## Aknowledgments

The authors would like to thank Dr. Nicholas Meyerson and Emma Rose Worden-Sapper for providing influenza virus, and Dr. Christopher Sullivan for comments on the manuscript.

## Author contributions

J.M.B. and R.P. conceived the project. J.M.B. performed influenza infections, poly(I:C) transfections, western blots, RT-qPCR, microscopy and ELISAs. A.R.G. performed DENV infections. J.M.B., A.R.G., S.L.S., and R.P. interpreted data. J.M.B. and R.P. wrote the manuscript.

## Data and materials availability

Correspondence and material request should be addressed to Dr. Roy Parker. All data needed to evaluate the conclusions in the paper are present in the paper and/or the Supplementary Materials. Key primary data such as raw microscopy images have been deposited in Mendeley data server (Burke, James (2021), “RNase L limits host and viral protein synthesis via inhibition of mRNA export”, Mendeley Data, V1, doi: 10.17632/xz46mfkg29.1 http://dx.doi.org/10.17632/xz46mfkg29.1). Additional primary data will be provided upon request.

## Supplemental figures and legends

**Fig. S1.**
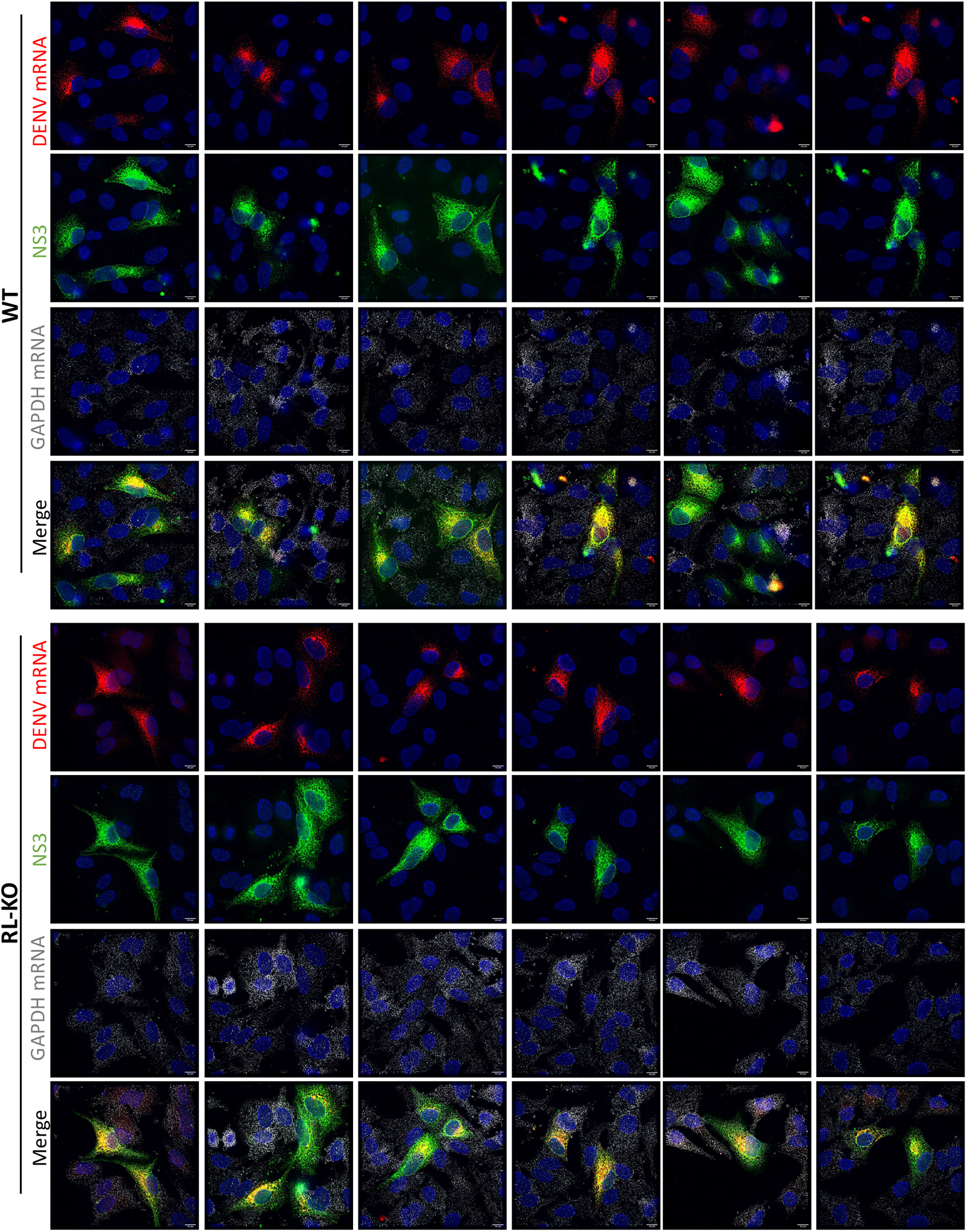
RNase L-mediated mRNA decay does not affect DENV mRNA or protein level. smFISH for *GAPDH* and *DENV* mRNAs and immunofluorescence detection of DENV NS3 protein in wild-type (WT) and RNase L-KO (RL-KO) A549 cells forty-eight hours post-infection with DENV (MOI=0.1).

**Fig. S2.**
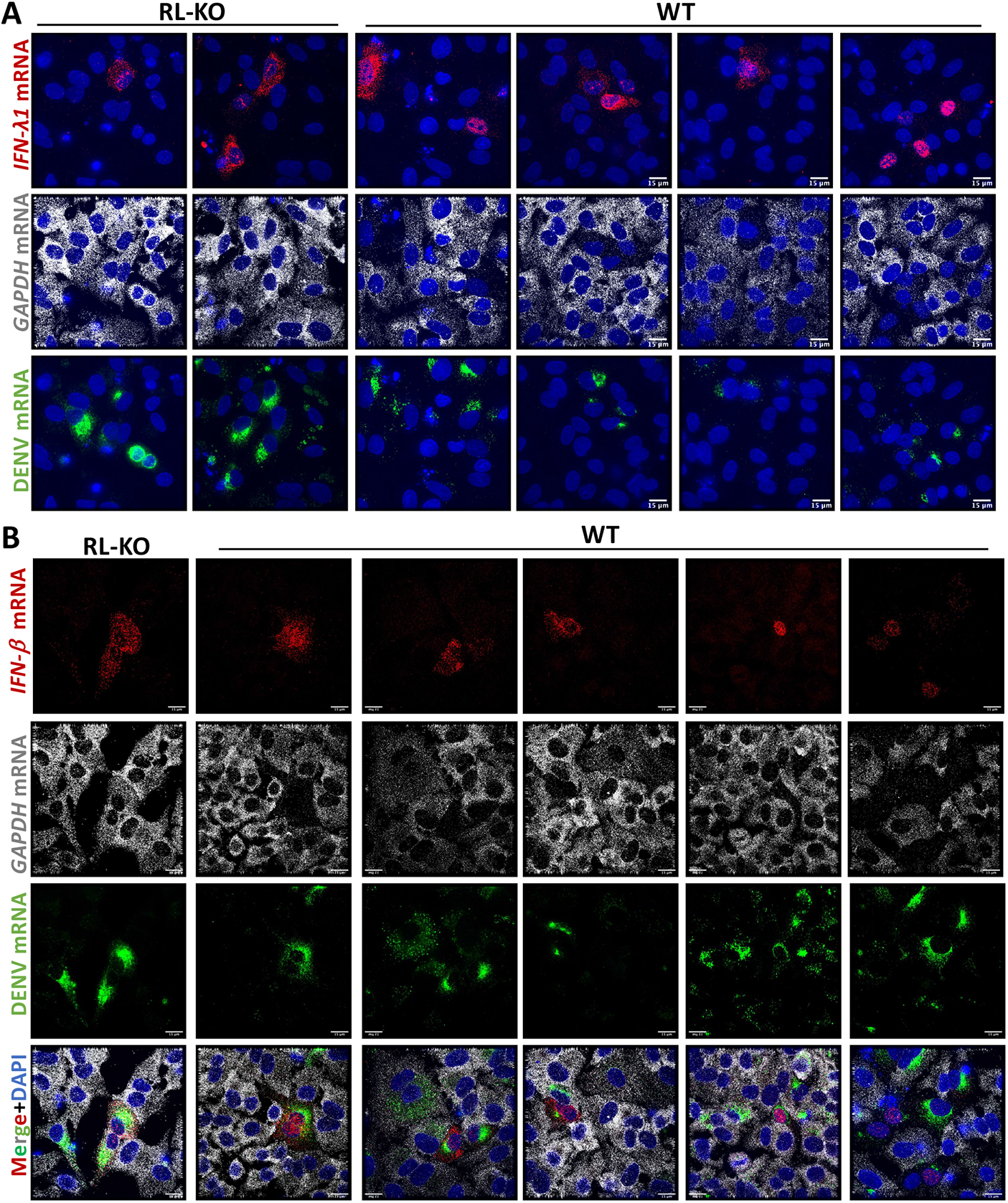
DENV activates RNase L-mediated mRNA decay escaped by antiviral mRNAs. (**A**) smFISH for *IFN-λ1*, *GAPDH,* and *DENV* mRNAs in wild-type (WT) and RNase L-KO (RL-KO) A549 cells forty-eight hours post-infection with DENV (MOI=0.1). (**B**) Similar to (**A**) but for *IFN-β*.

**Fig. S3.**
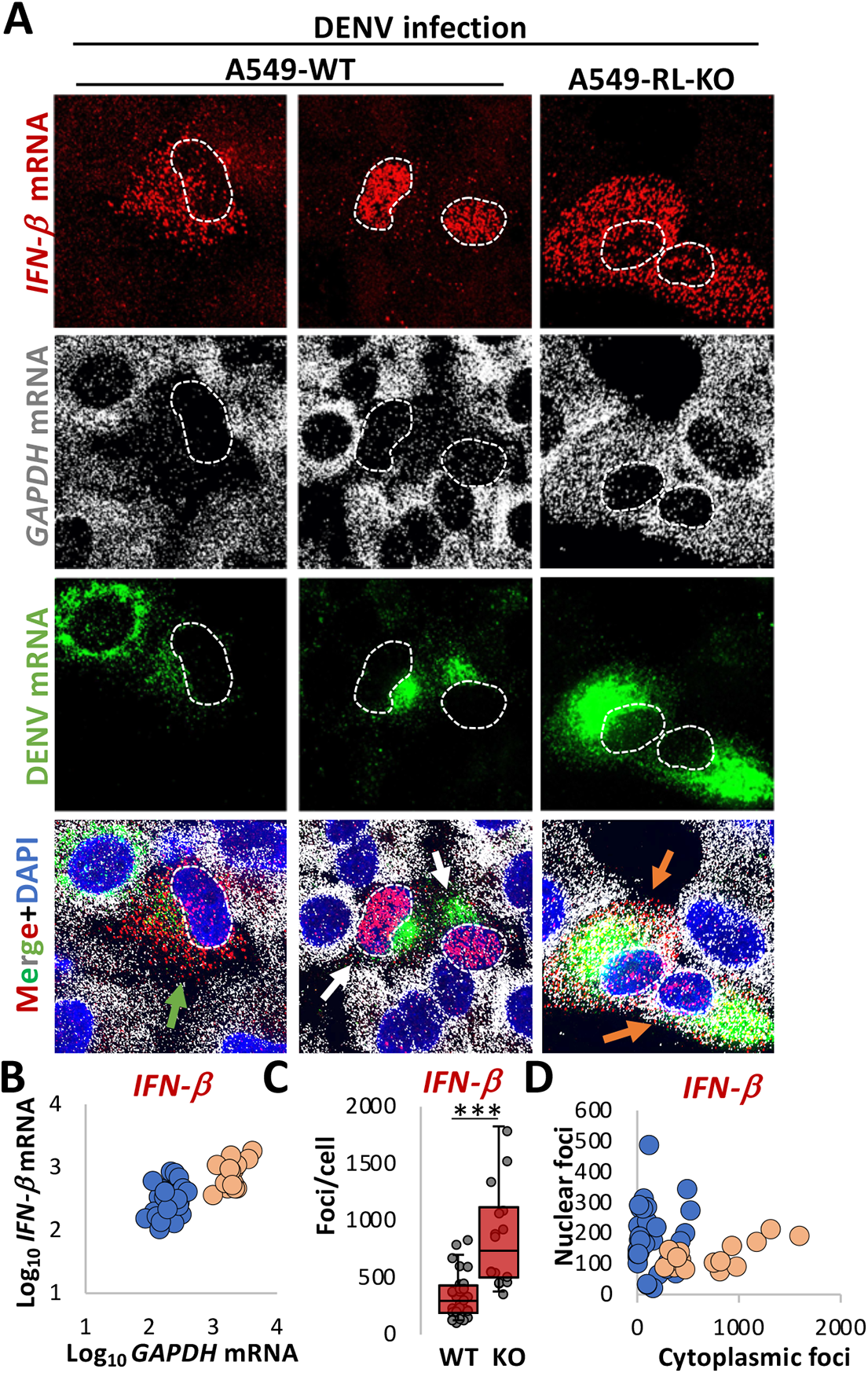
RNase L activation inhibits *IFN-β* mRNA export in response to DENV infection. (**A**) smFISH for *IFN-β* mRNA, *GAPDH* mRNA, and *DENV* mRNA in WT and RL-KO A549 cells forty-eight hours post-infection with DENV (MOI=0.1). Orange arrow: DENV-positive but RNase L not activated (*GAPDH* mRNA not degraded); green arrow: DENV positive, RNase L activated (*GAPDH* mRNA degraded), and *IFN-β* mRNA localized to cytoplasm; white arrow: DENV-positive, RNase L activated, and IFN mRNAs retained in the nucleus. (**B**) Scatter plot of *GAPDH* mRNA and *IFN-β* mRNA in individual cells. (**C**) Quantification of smFISH foci in WT and RL-KO cells infected with DENV and that induced IFN-*β* mRNA expression. (**D**) Scatter plot of the number of smFISH foci of *IFN-β* mRNA localized to the nucleus or cytoplasm in individual WT and RL-KO cells as represented in (**A**).

**Fig. S4.**
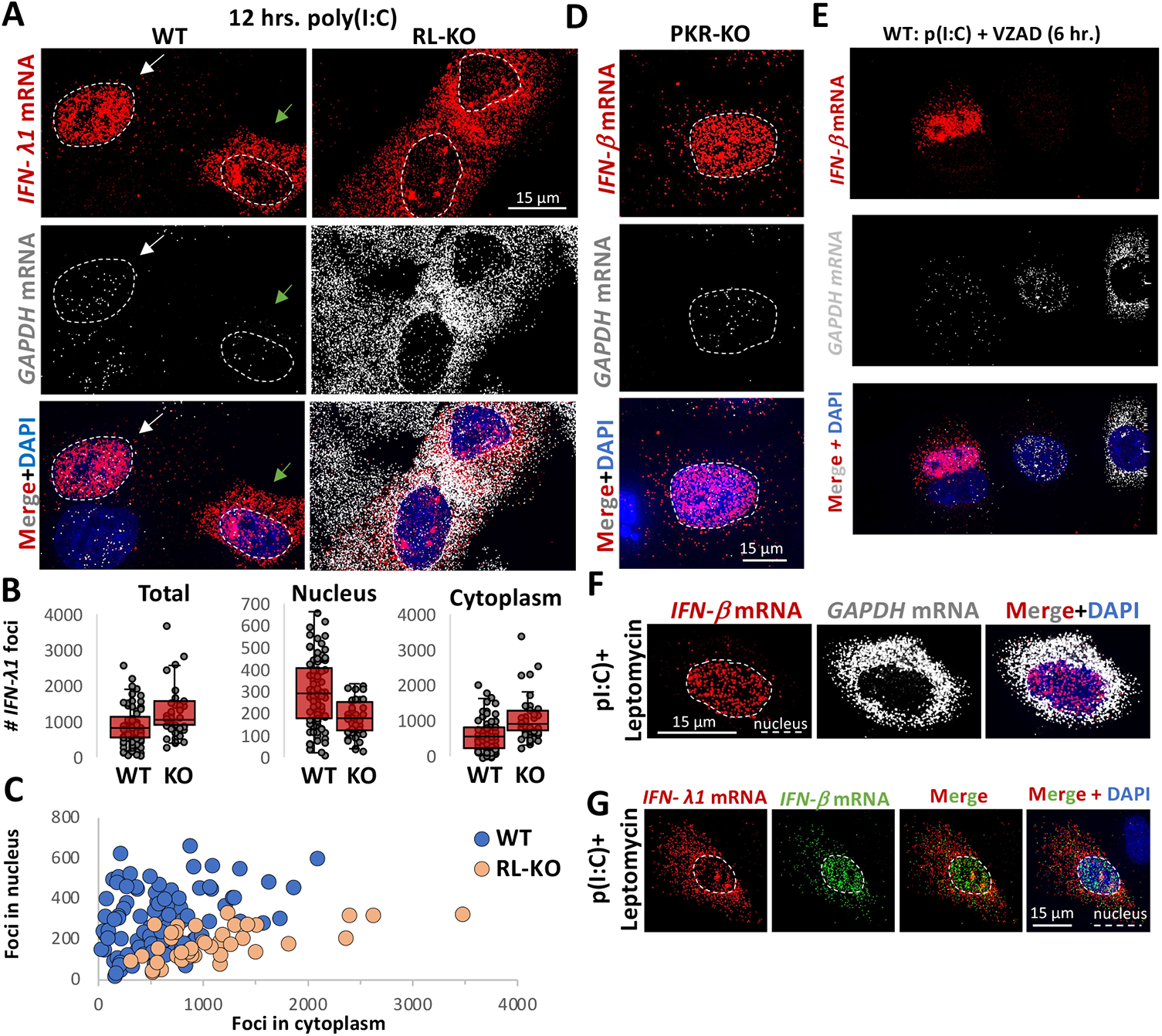
RNase L activation inhibits *IFN* mRNA export in response to dsRNA. (**A**) smFISH for *IFN-λ1* and *GAPDH* mRNAs in wild-type (WT) and RNase L-KO (RL-KO) A549 cells eight hours post-poly(I:C) lipofection. (**B**) quantification of *IFN-λ1* smFISH foci from WT and RL-KO cells eight-hours post-poly(I:C), as represented in (**C**). Each dot represents *IFN-β* levels from an induvial cell. (**C**) scatter plot of the nuclear (y-axis) and cytoplasmic (x-axis) levels of *IFN-β* smFISH in individual. smFISH for *IFN-β* and *GAPDH* mRNAs in PKR-KO A549 cells eight hours post-poly(I:C) lipofection.**(D)** smFISH for *IFN-β* mRNA in WT A549 cells six hours post-poly(I:C) transfection. Cells were treated with VZAD pan caspase inhibitor 5 minutes before transfection. (**F**) smFISH for *IFN-β* and *GAPDH* mRNAs in RL-KO cells six hours post-poly(I:C) lipofection and treated with leptomycin two hours post-transfection. (**G**) smFISH for *IFN-β* and *IFN-λ1* mRNAs in RL-KO cells six hours post-poly(I:C) lipofection and treated with leptomycin two hours post-transfection.

**Fig. S5.**
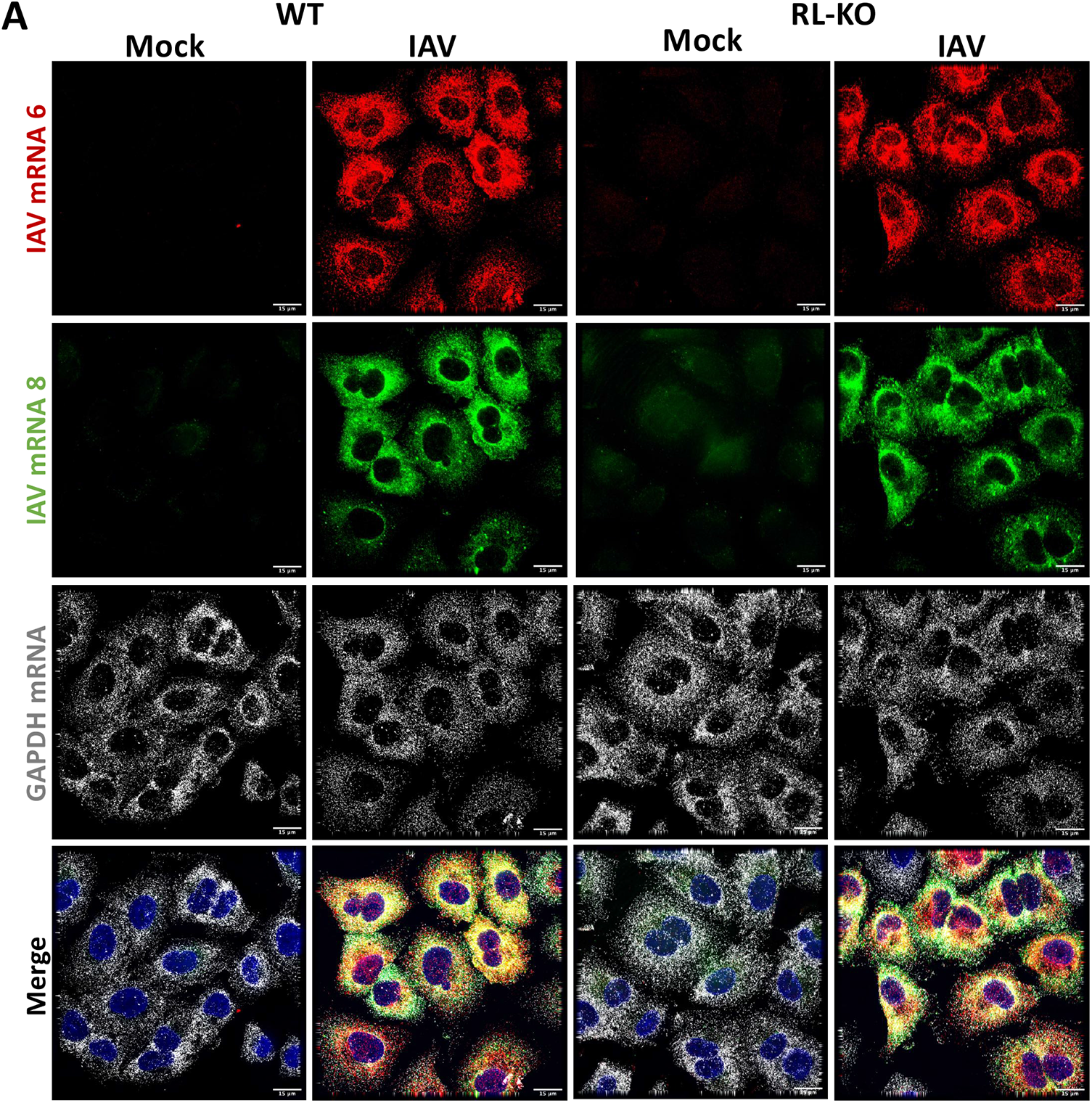
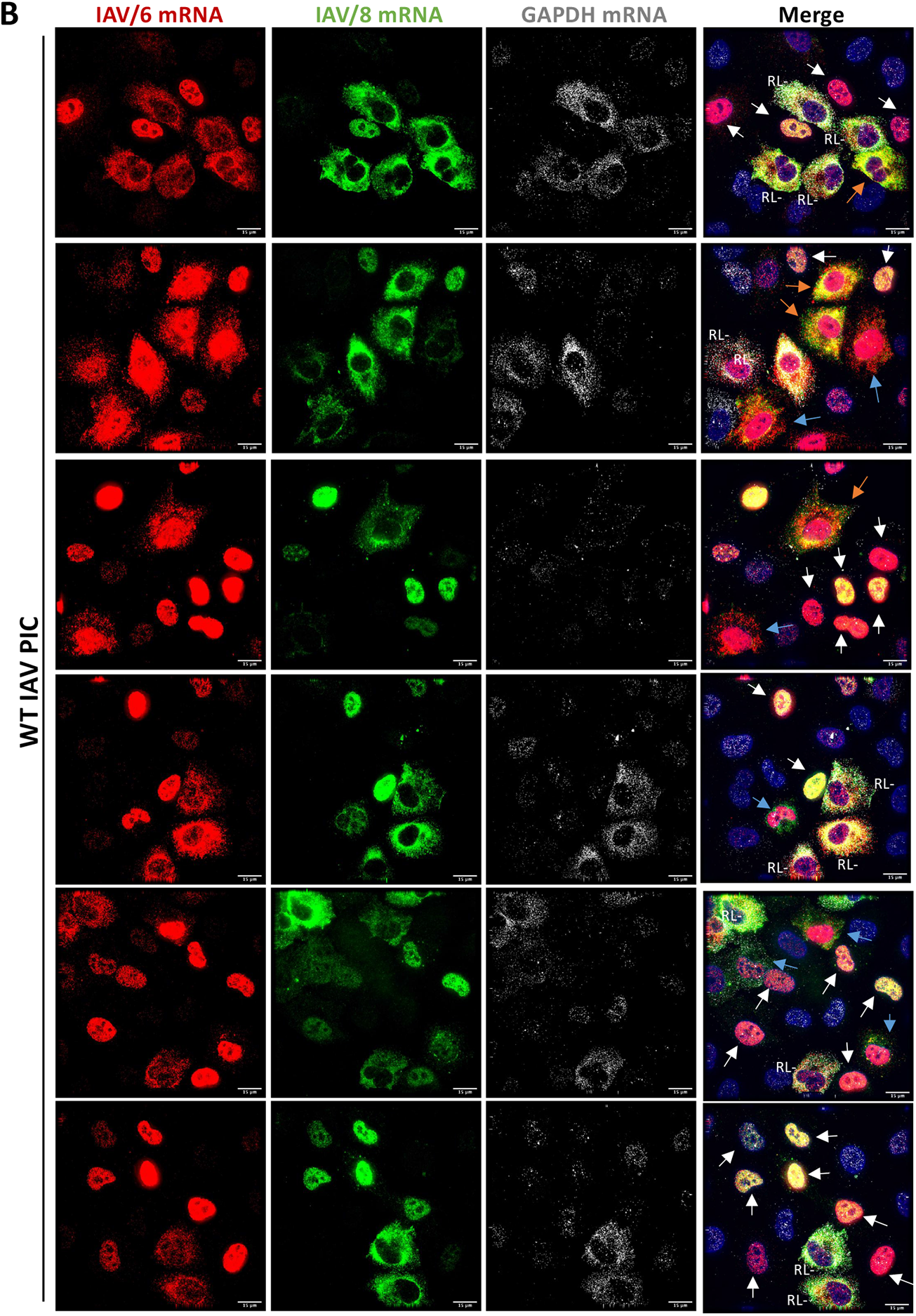
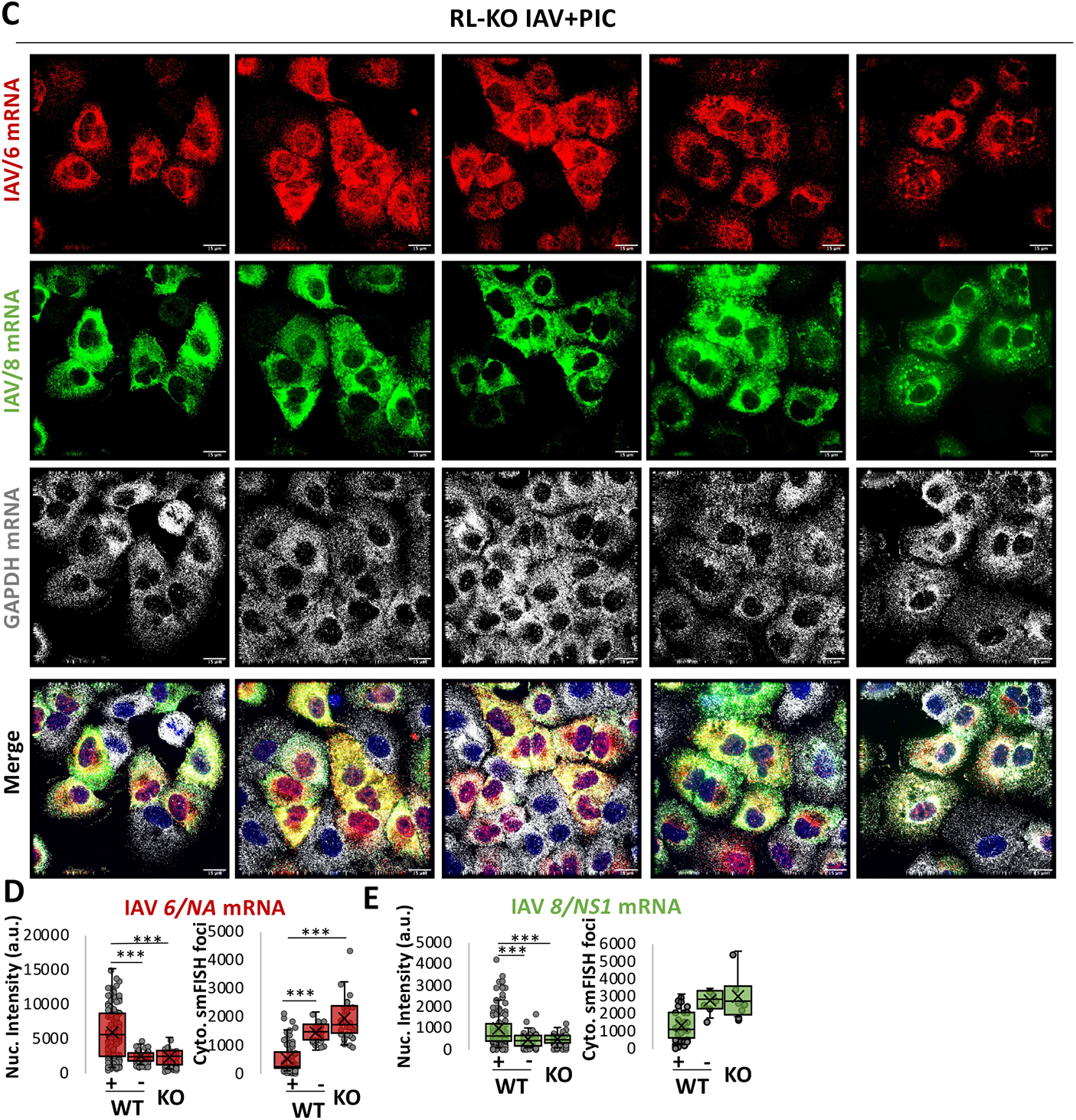
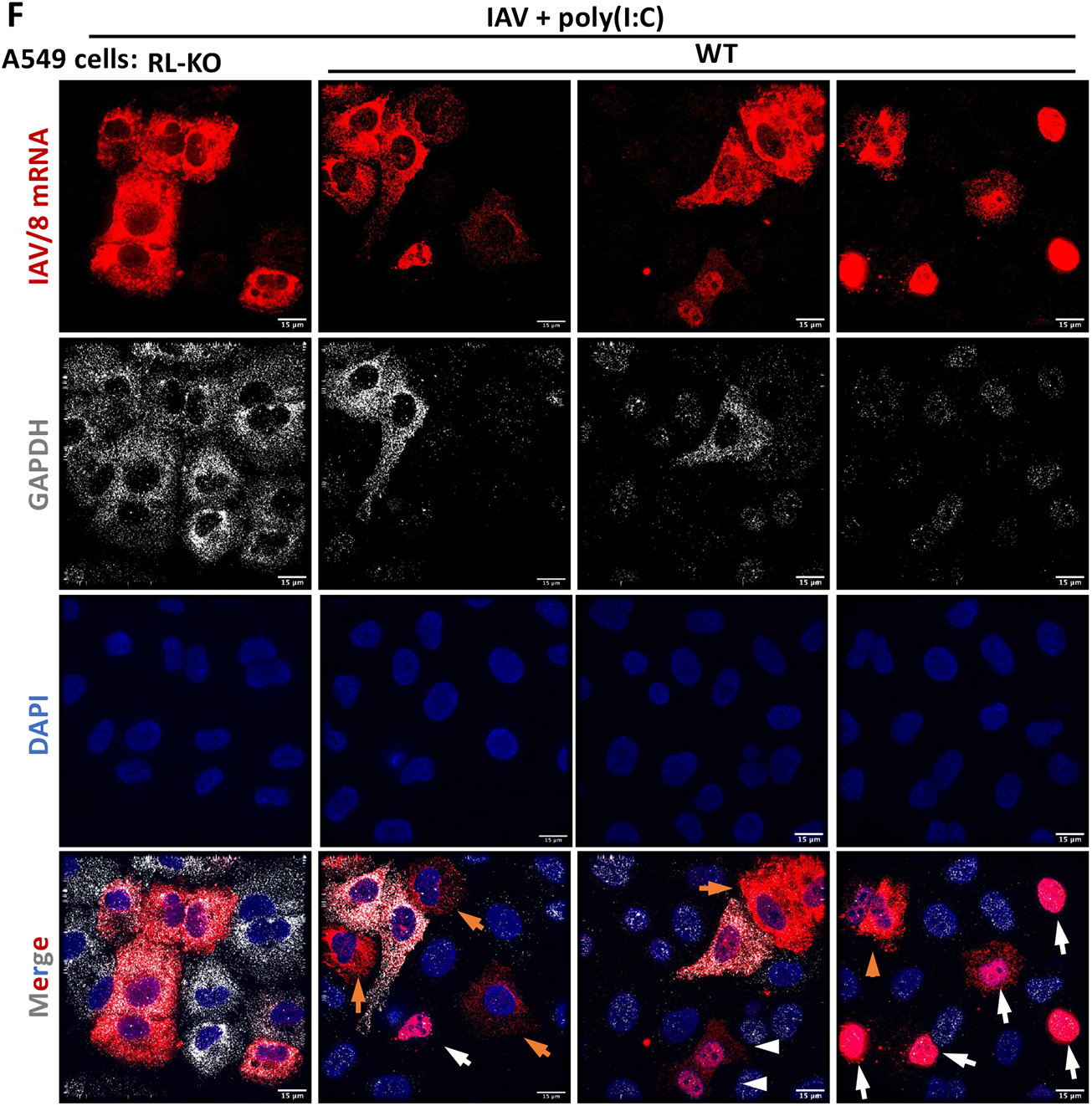
RNase L-mediated inhibition of influenza virus mRNA export. (**A**) smFISH for IAV *6/NA* and *8/NS1* mRNAs and *GAPDH* mRNA seven hours post-infection with influenza A/Udorn/72 virus (MOI=0.5). Mock infection was also performed. We note that substantial *GAPDH* mRNA decay was not observed in infected cells, indicating a lack of RNase L activation, which is consistent with NS1-mediated inhibition of RNase L activation as previously described. We also note that IAV mRNAs are predominantly cytoplasmic, indicating their efficient export. (**B** and **C**) smFISH for IAV *6/NA* and *8/NS1* mRNAs and *GAPDH* mRNA seven hours post-infection of WT (**B**) or RL-KO (**C**) A549 cells with influenza A/Udorn/72 virus (MOI=0.5). Cells were transfected with poly(I:C) one-hour post-poly(I:C) to promote RNase L activation, which can be observed by the loss of *GAPDH* mRNA. Multiple fields are shown to illustrate the broad and diverse phenotypes of IAV *6/NA* and *8/NS1* mRNA localization. (**D** and **E**) Quantification of nuclear intensity (left graph) and cytoplasmic smFISH foci (right graph) of IAV *6/NA* mRNA (**D**) or IAV *8/NS1* mRNA (**E**) from WT cells, in which RNase L was activated (RL+) or not activated (RL-), and RL-KO cells. (**F**) similar to (**A** and **B**) but for *8/NS1* mRNAs and *GAPDH* mRNA to better illustrate escape of *8/NS1* mRNA from RNase L-mediated mRNA decay, which can be observed in cells that degrade *GAPDH* mRNA but not IAV *8/NS1* mRNA (orange arrows).

**Fig. S6.**
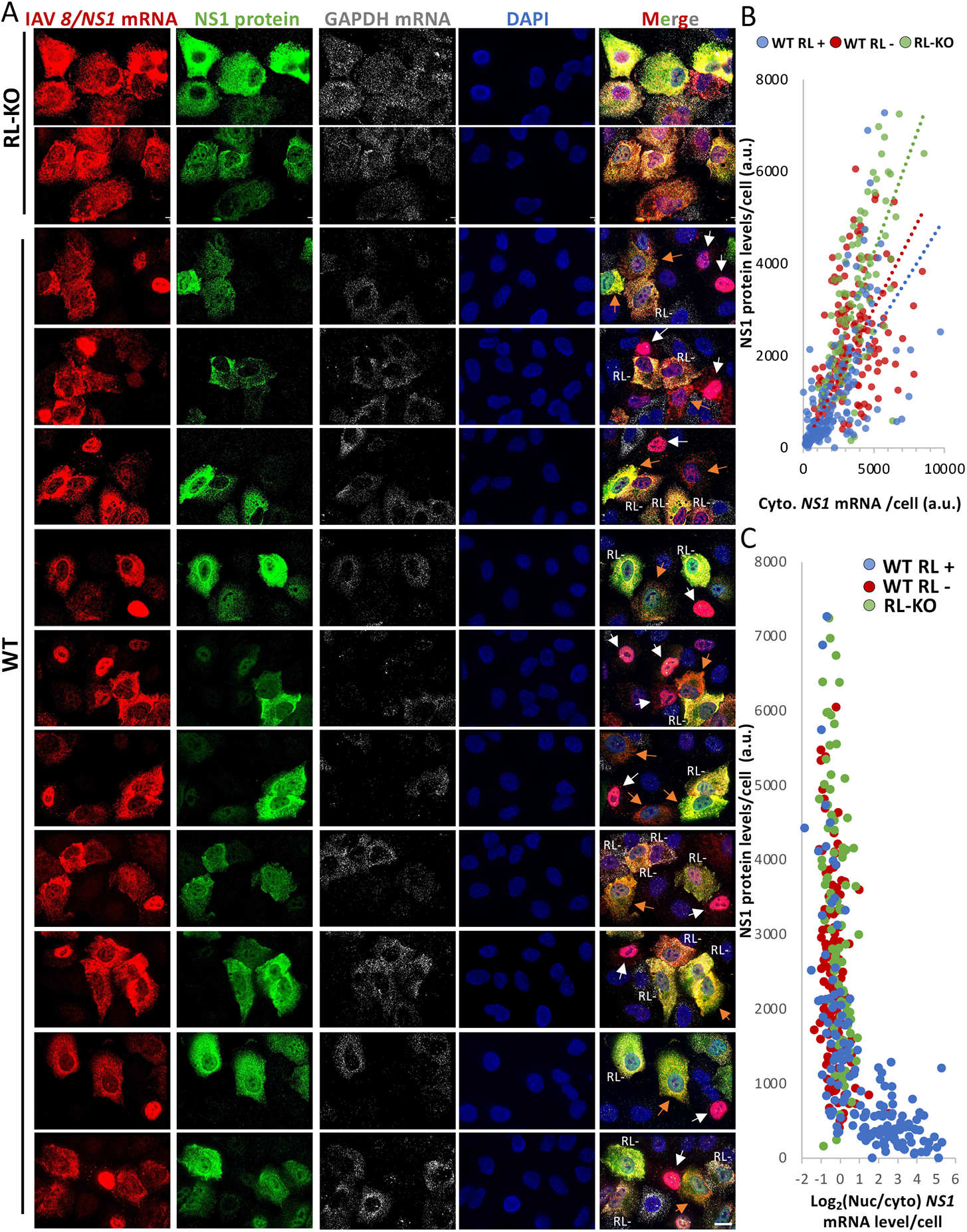
RNase L-mediated inhibition of IAV mRNA export is necessary to inhibit IAV protein synthesis. (**A**) smFISH for IAV *8/NS1* mRNA and *GAPDH* mRNAs and immunofluorescence of NS1 protein in WT and RL-KO A549 cells seven-hours post-infection with influenza A/Udorn.72 virus (MOI=0.5). Cells were transfected with poly(I:C) one-hour post-infection to activate RNase L. Two fields of view are shown for RL-KO cells, and ten fields of view for WT cells to display the diversity in IAV mRNA localization and *NS1* levels in WT cells. (**B**) Scatter plot of *NS1* protein intensity and cytoplasmic IAV 8/NS1 mRNA intensity from WT cells that activated RNase L (RL+) or did not activate RNase L (RL-) and RL-KO cells as represented in (**A**) shows that increased cytoplasmic IAV 8/NS1 mRNA levels correlate with increased *NS1* protein intensity in both WT (RL- and RL+) and RL-KO cells. (**C**) scatter plot of *NS1* protein intensity (y-axis) and the ratio of nuclear/cytosolic *8/NS1 mRNA* intensity (x-axis) from WT cells that active RNase L (RL+) or did not activate RNase L (RL-) and RL-KO cells as represented in (**A**).

**Fig. S7.**
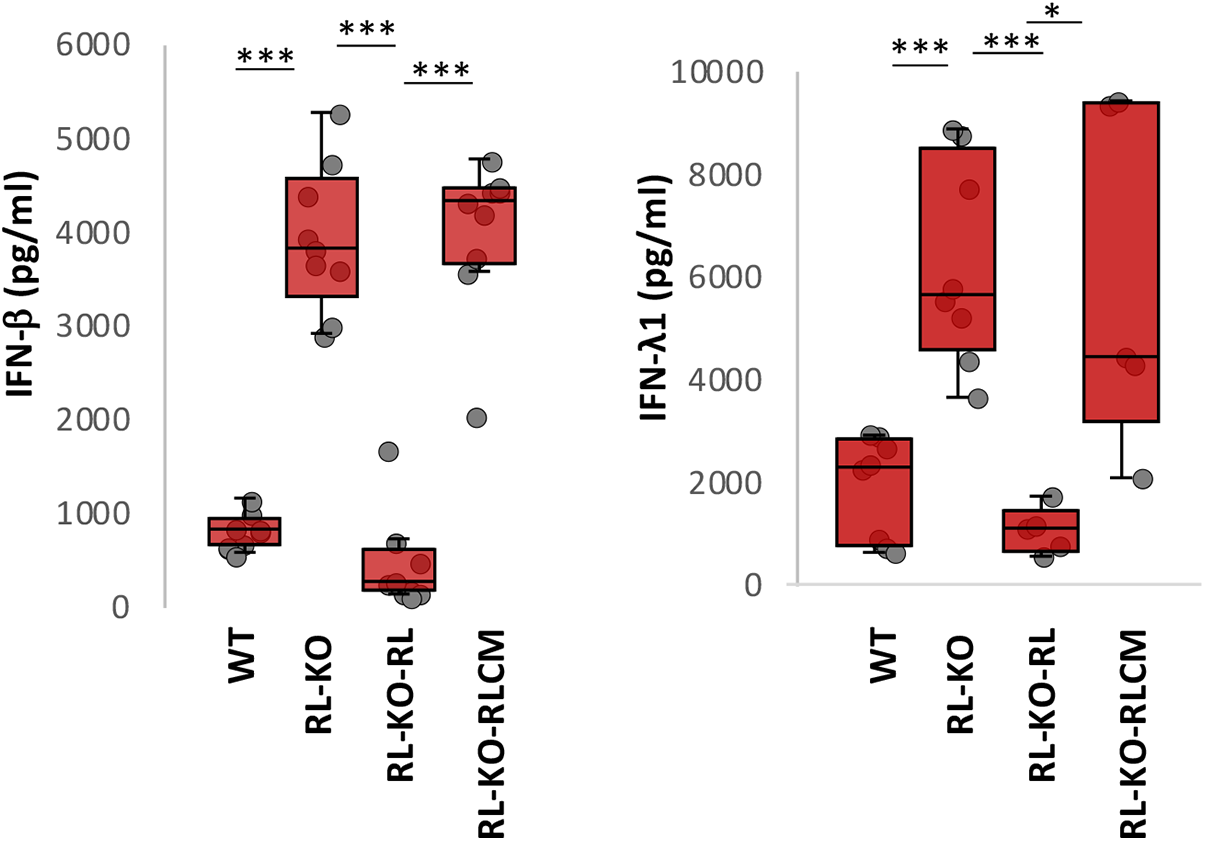
RNase L catalytic activity inhibits type I and type III interferon production. ELISA for IFN-β and IFN-λ1 from WT, RL-KO, and RL-KO cells stably expressing either RNase L (RL) or RNase L-R667A catalytic mutant (RL-CM) sixteen hours post-poly(I:C) lipofection.

**Fig. S8.**
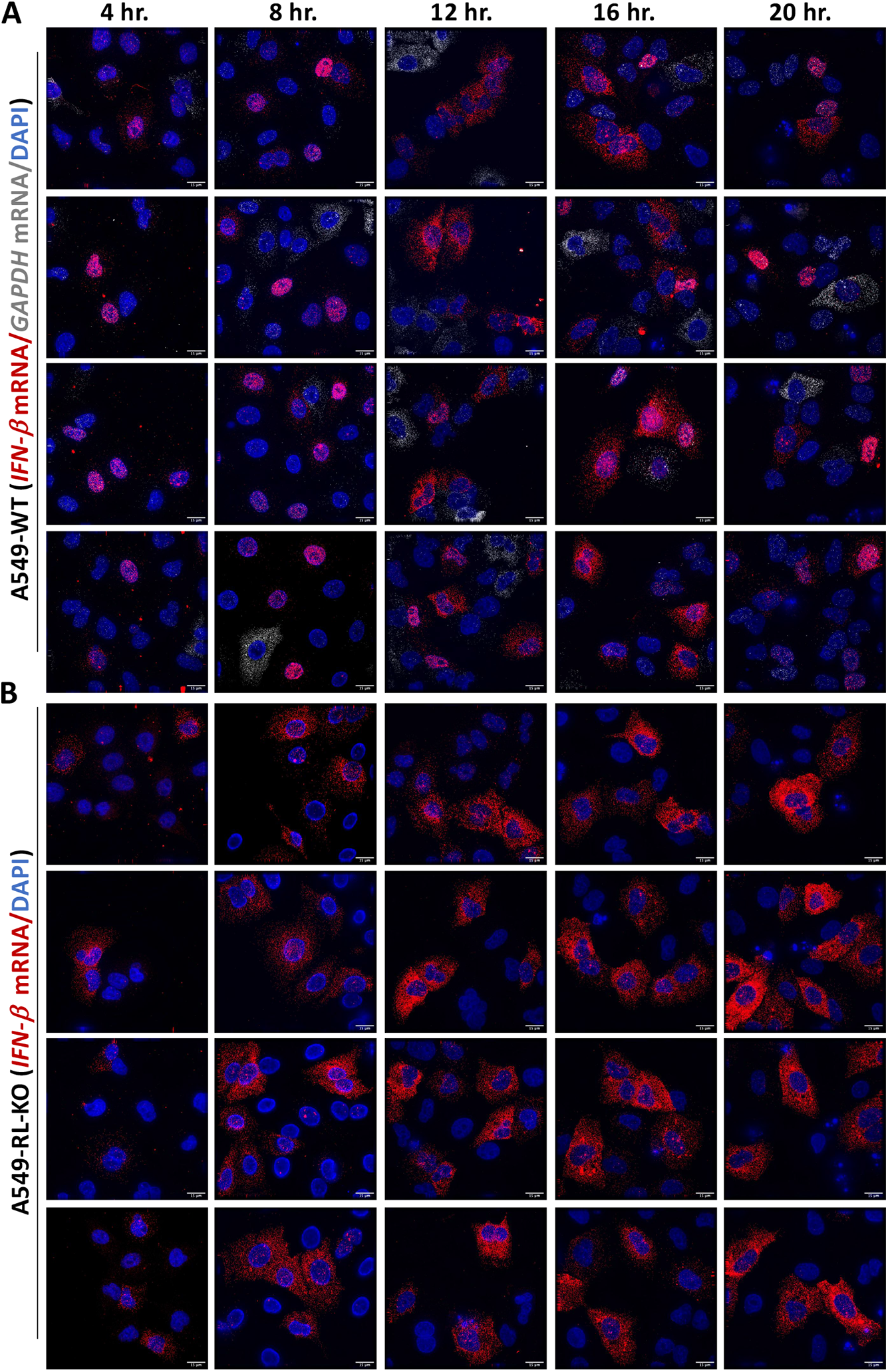
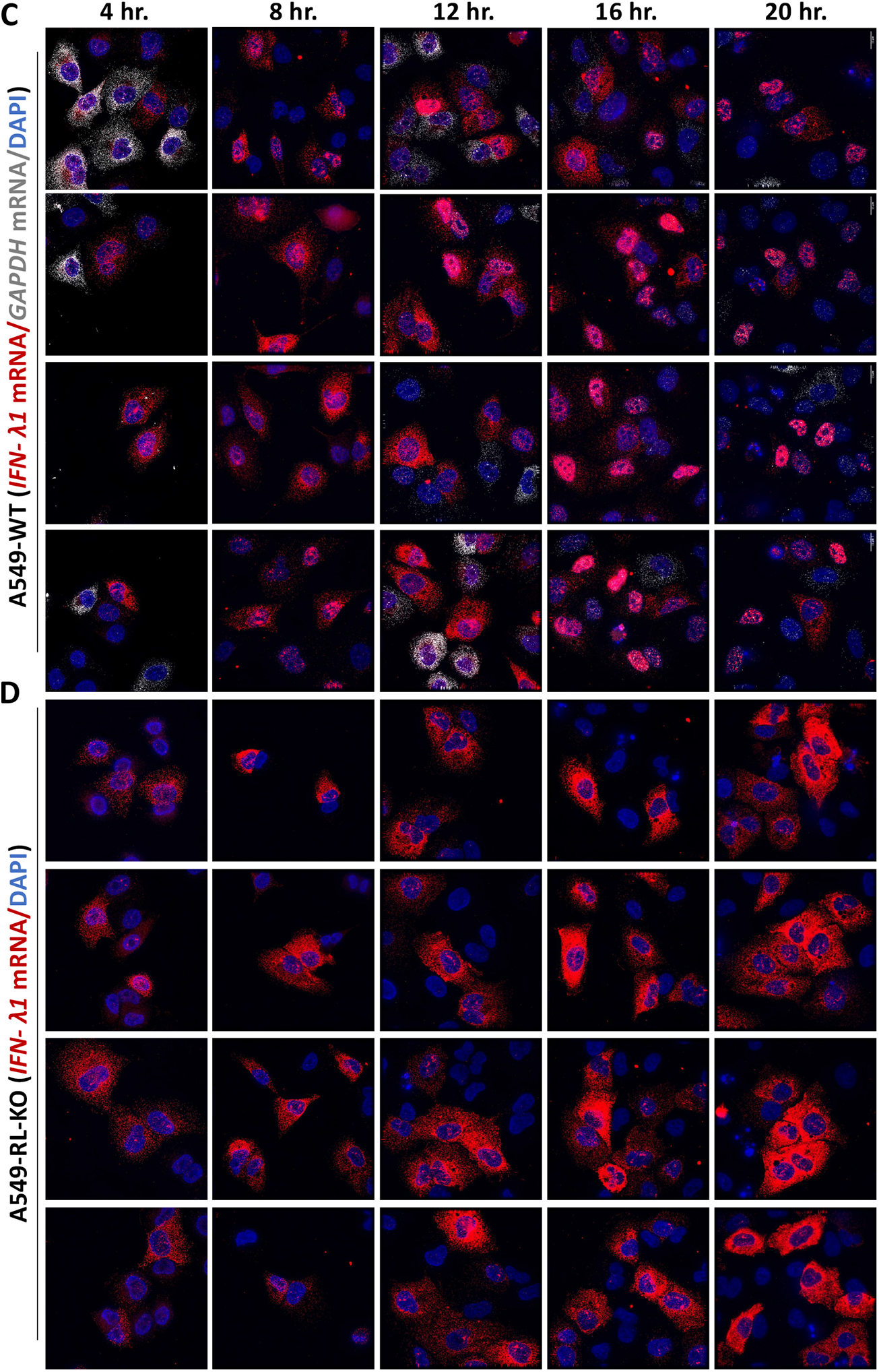
IFN mRNA localization over time in WT and RL-KO cells. (**A**) Four representative images of smFISH for *IFN-*β and *GAPDH* mRNA in WT A549 cells at indicated times post-poly(I:C) lipofection. Nuclei were stained with DAPI. (**B**) smFISH for *IFN-*β in RL-KO A549 cells at indicated times post-poly(I:C) lipofection. *GAPDH* mRNA smFISH is not shown to visualize cytoplasmic *IFN-*β mRNA due to space constraints. (**C** and **D**) similar to (**A** and **B**) but for *IFN-λ1* mRNA.

**Fig. S9.**
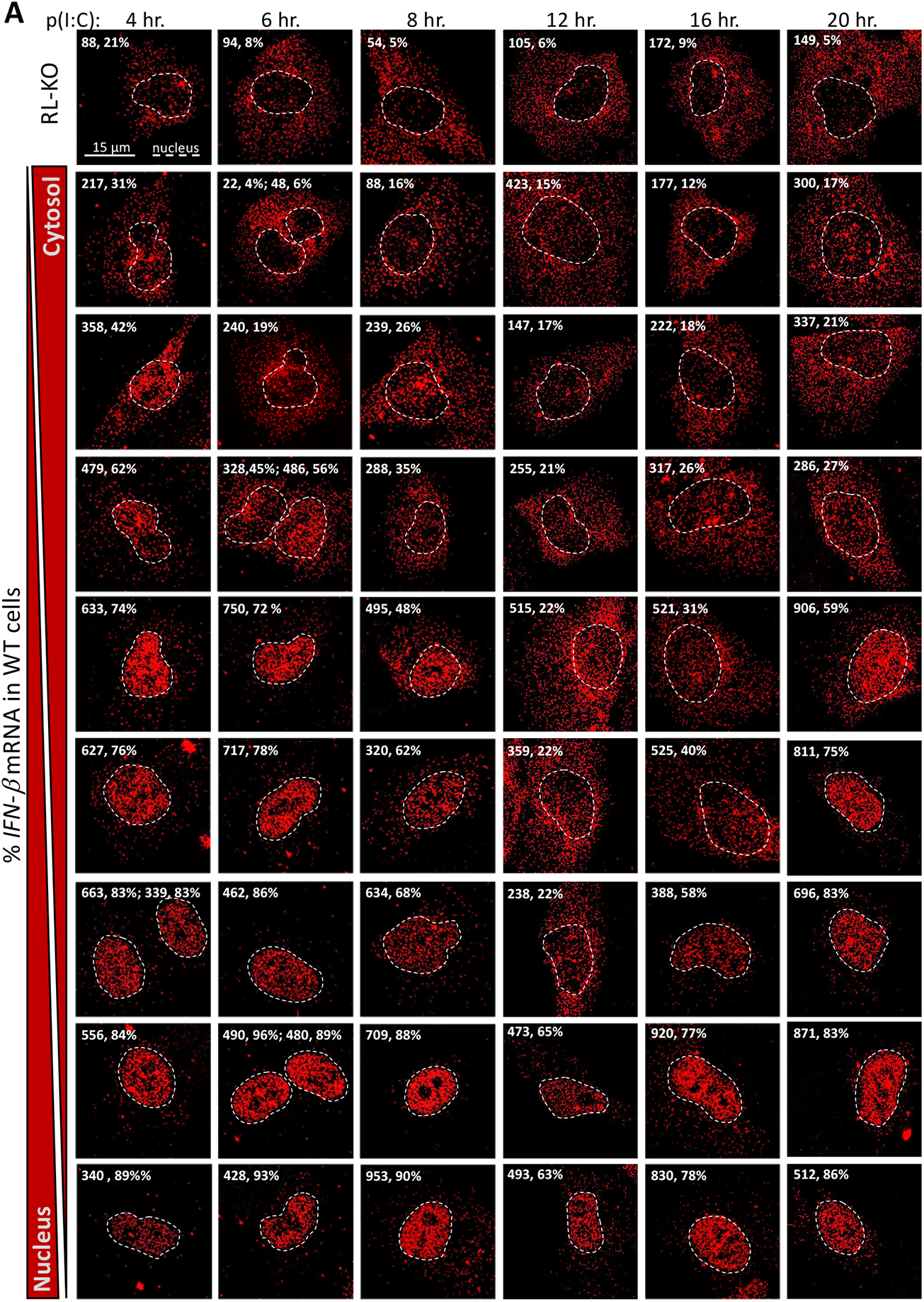
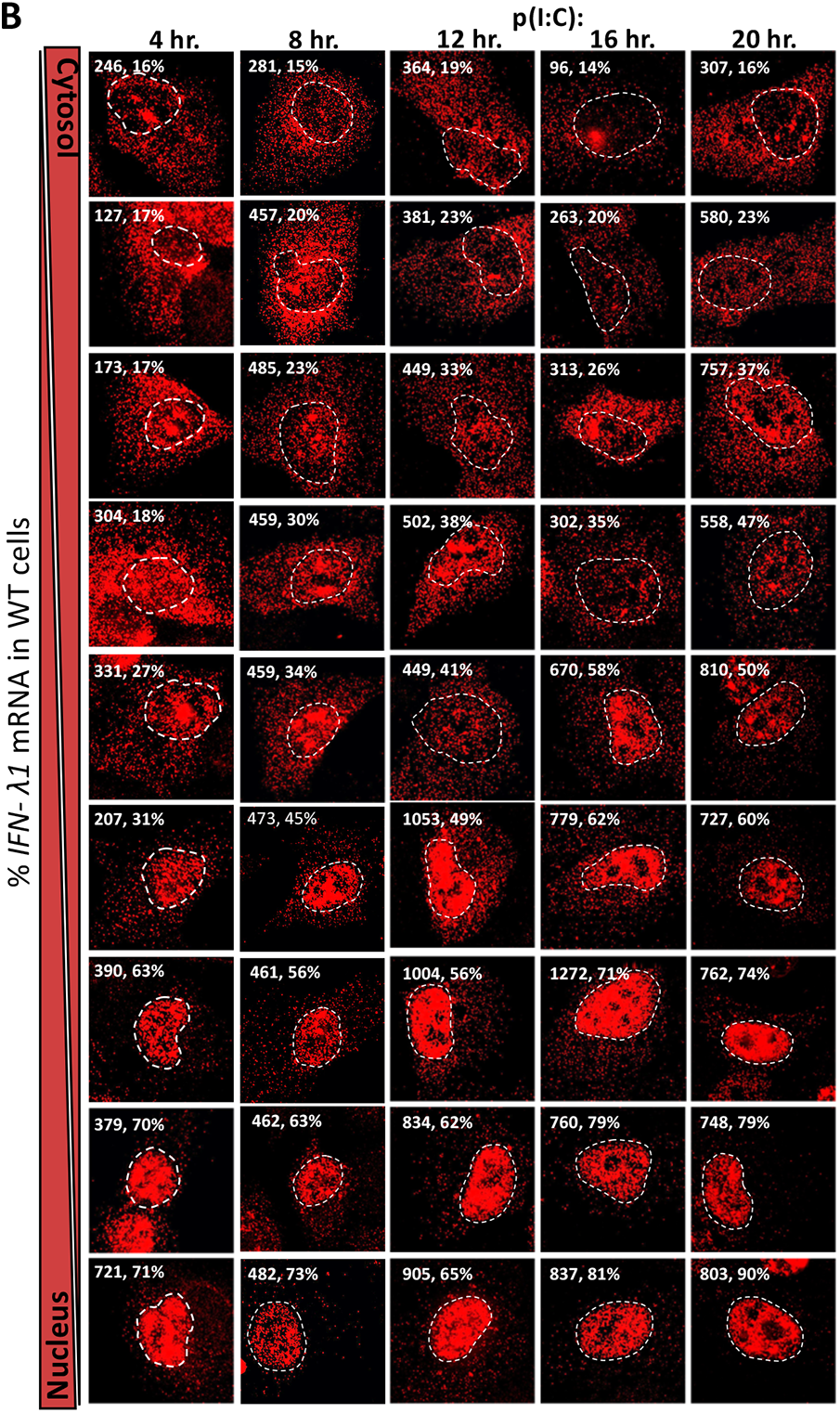
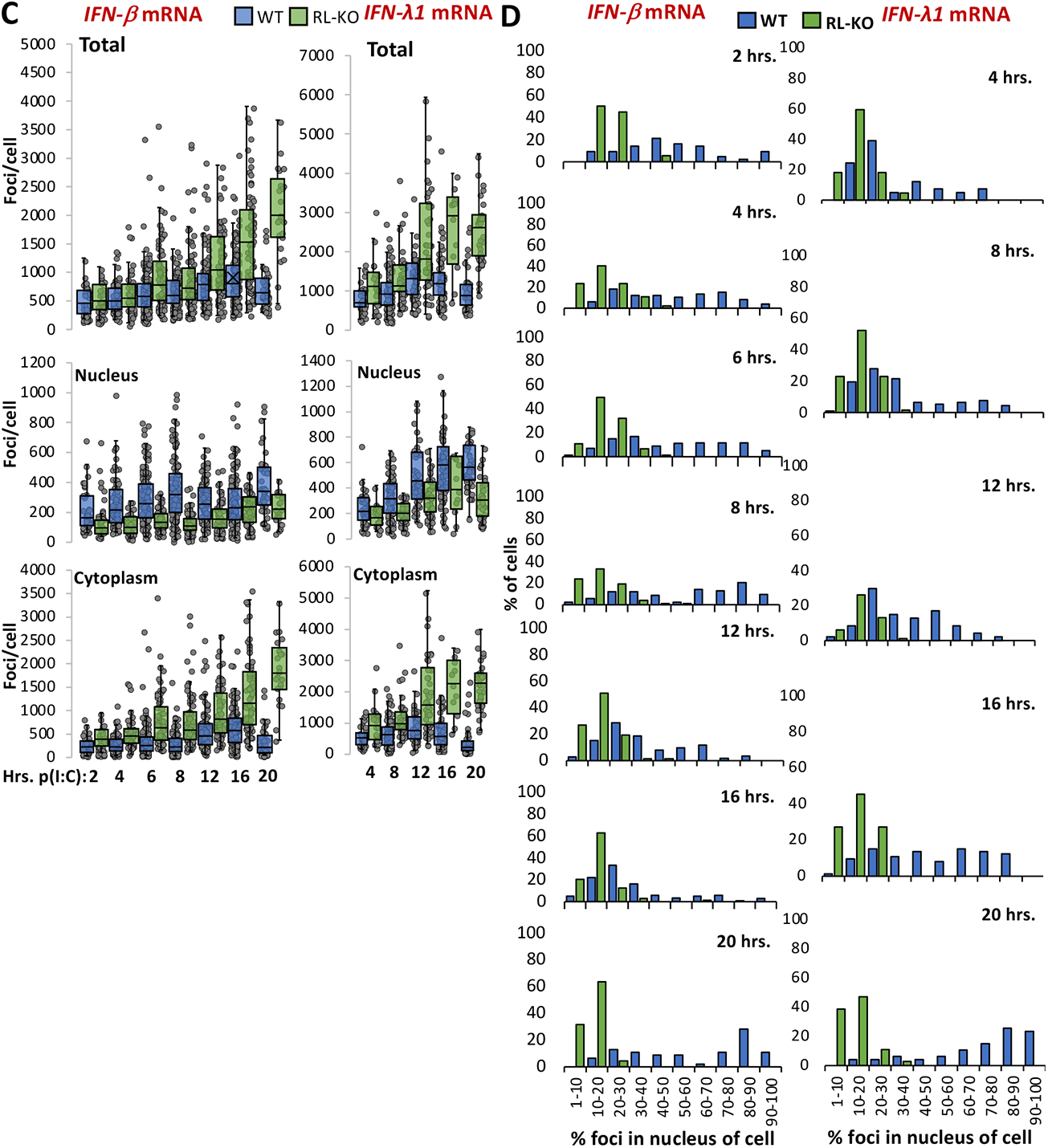
Quantification of RNase L-mediated nuclear retention of antiviral mRNA. (**A** and **B**) Representative WT A549 cells displaying varying degrees of nuclear retention of *IFN-β* (**A**) or *IFN-λ1* (**B**) mRNA post-poly(I:C). The number of nuclear smFISH foci (left) and the percentage of smFISH foci localized to the nucleus (right) are shown in the top right corners. Nuclei were determined by DAPI staining (not shown for space) and are outlined. (**C**) Quantification of total, nuclear, or cytoplasmic *IFN-β* or *IFN-λ1* smFISH foci in WT and RL-KO A549 cells at indicated time post-poly(I:C), as represented in (**A**) and (**B**). Each dot represents the number of foci in an individual cell. (**D**) Histograms showing the percentage of cells displaying binned ranges of *IFN-β* and *IFN-λ1* localization to the nucleus.

